# High-throughput screens of PAM-flexible Cas9 variants for gene knock-out and transcriptional modulation

**DOI:** 10.1101/2020.01.22.916064

**Authors:** Mateusz Legut, Zharko Daniloski, Xinhe Xue, Dayna McKenzie, Xinyi Guo, Hans-Hermann Wessels, Neville E. Sanjana

## Abstract

A key limitation of the commonly-used CRISPR enzyme *S. pyogenes* Cas9 is the strict requirement of an NGG protospacer-adjacent motif (PAM) at the target site, which reduces the number of accessible genomic loci. This constraint can be limiting for genome editing applications that require precise Cas9 positioning. Recently, two Cas9 variants with a relaxed PAM requirement (NG) have been developed (xCas9 and Cas9-NG) but their activity has been measured at only a small number of endogenous sites. Here we devised a high-throughput Cas9 pooled competition screen to compare the performance of both PAM-flexible Cas9 variants and wild-type Cas9 at thousands of genomic loci and across 3 modalities (gene knock-out, transcriptional activation and suppression). We show that PAM flexibility comes at a substantial cost of decreased DNA targeting and cutting. Of the PAM-flexible variants, we found that Cas9-NG outperforms xCas9 regardless of genome engineering modality or PAM. Finally, we combined xCas9 mutations with those of Cas9-NG, creating a stronger transcriptional modulator than existing PAM-flexible Cas9 variants.

## Introduction

Type II CRISPR-Cas9 enzymes are RNA-programmable endonucleases that have been used in diverse DNA-targeting applications, including gene knock-out and knock-in, mutagenesis, gene activation and inhibition, base editing, and CpG methylation (Jinek et al., 2012). Cas9 enzymes, including the most commonly used *S. pyogenes* Cas9 (Cas9), recognize target DNA sequences that are complementary to their guide RNA spacer and that contain a protospacer adjacent motif (PAM). Although mismatches between the target DNA and portion of the guide RNA can be tolerated, the presence of the PAM is a strict requirement, which imposes a limit on the number of targetable genomic loci (Hsu et al., 2013). While the availability of PAM sites (such as NGG for Cas9) is typically not a problem for CRISPR-mediated gene knockout because nearly all proteincoding exons can be targeted (Meier et al., 2017), the optimal targeting space for transcriptional modulation (inhibition or activation) is usually smaller, between 50 and 100 nucleotides (Sanson et al., 2018). Other common genome editing tasks, such as homology-directed repair or base editing, require an even narrower window for Cas9 positioning, with the desired target site placed at a precise position from the PAM sequence (e.g. 10 - 20 nucleotides for homology-directed repair or 13 - 17 nucleotides for base editing) (Findlay et al., 2014; Komor et al., 2016).

To address this problem, several Cas9 orthologs and other CRISPR nucleases from different bacterial species have been characterized, such as *S. aureus* Cas9 and Cas12a/Cpf1 (Kim et al., 2016; Ran et al., 2015). However, none of them have a simpler PAM requirement than Cas9. Initial attempts at developing more PAM-flexible Cas9 variants through structure-based design or directed evolution yielded enzymes recognizing slightly altered PAMs but still requiring a three-nucleotide motif (Kleinstiver et al., 2015). Recently, two Cas9 variants capable of recognizing an NG PAM were generated, one through phage-assisted continuous evolution (xCas9), and the other through structure-guided design (Cas9-NG) (Hu et al., 2018; Nishimasu et al., 2018). These Cas9 variants were characterized primarily in terms of their nuclease activity at several endogenous genomic loci, and their relative performance at NG sites was highly variable. One of these PAM-flexible variants, xCas9, led to superior CRISPR activation (CRISPRa) when fused to VP64-p65-Rta (VPR) over wild-type dCas9-VPR, with higher transcriptional activation for all sgRNAs tested. This is presumably due to the directed evolution selection pressure — transcriptional activation and not nuclease activity — used to derive xCas9.

Given the utility of PAM-flexible Cas9 enzymes for precise genome engineering, we designed an unbiased, massively-parallel competition assay to compare Cas9 enzyme variants at thousands of target sites in the human genome. We benchmarked both PAM-flexible enzymes head-to-head with Cas9 for nuclease-driven loss-of-function, gene activation and gene repression. Across all 3 modalities, we found that PAM flexibility comes at the cost of markedly lower activity. Wild-type Cas9 outperformed both PAM-flexible variants at NGG sites for every modality tested. At NGH PAMs (H = A, C or T), we found that Cas9-NG is universally better than xCas9 and that xCas9 is often indistinguishable from the wild-type enzyme. We were able to partially rescue xCas9 nuclease activity by adding Cas9-NG mutations to create a new Cas9 variant, xCas9-NG. For gene activation, we found that xCas9-NG outperforms both xCas9 and Cas9-NG at both NGG and NGH PAMs. We expect that this novel PAM-flexible Cas9 will be useful for a multitude of genomeengineering applications where precise Cas9 positioning is required.

## Results

### A high-throughput competition screen to compare PAM-flexible Cas9 variants

To compare Cas9 variants across different PAM sites and different genome engineering tasks, we designed a high-throughput competition assay to test three Cas9 variants (wild-type [WT] Cas9, Cas9-NG and xCas9) and three different genetic perturbations (nuclease, transcriptional activation, and transcriptional repression) at thousands of target sites in the human genome (**Figure 1A**). For transcriptional activation (CRISPRa), we used nuclease-null versions of each Cas9 variant (D10A/H840A) fused to VPR proteins. VPR and other synergistic activators with multiple activation domains, such as SAM and SunTag, outperform single domain activators (Chavez et al., 2016). For transcriptional repression (CRISPR inhibition, CRISPRi), we tethered the nuclease-null variants to the KRAB repressor domain (Kearns et al., 2014). All Cas9 variant mutations were made on the same background using a human codon-optimized WT Cas9 from lentiCRISPRv2 (Sanjana et al., 2014) (**Figure S1A**) and we noticed no differences in protein expression between Cas9 variants (**Figure S2**).

**Figure 1.**
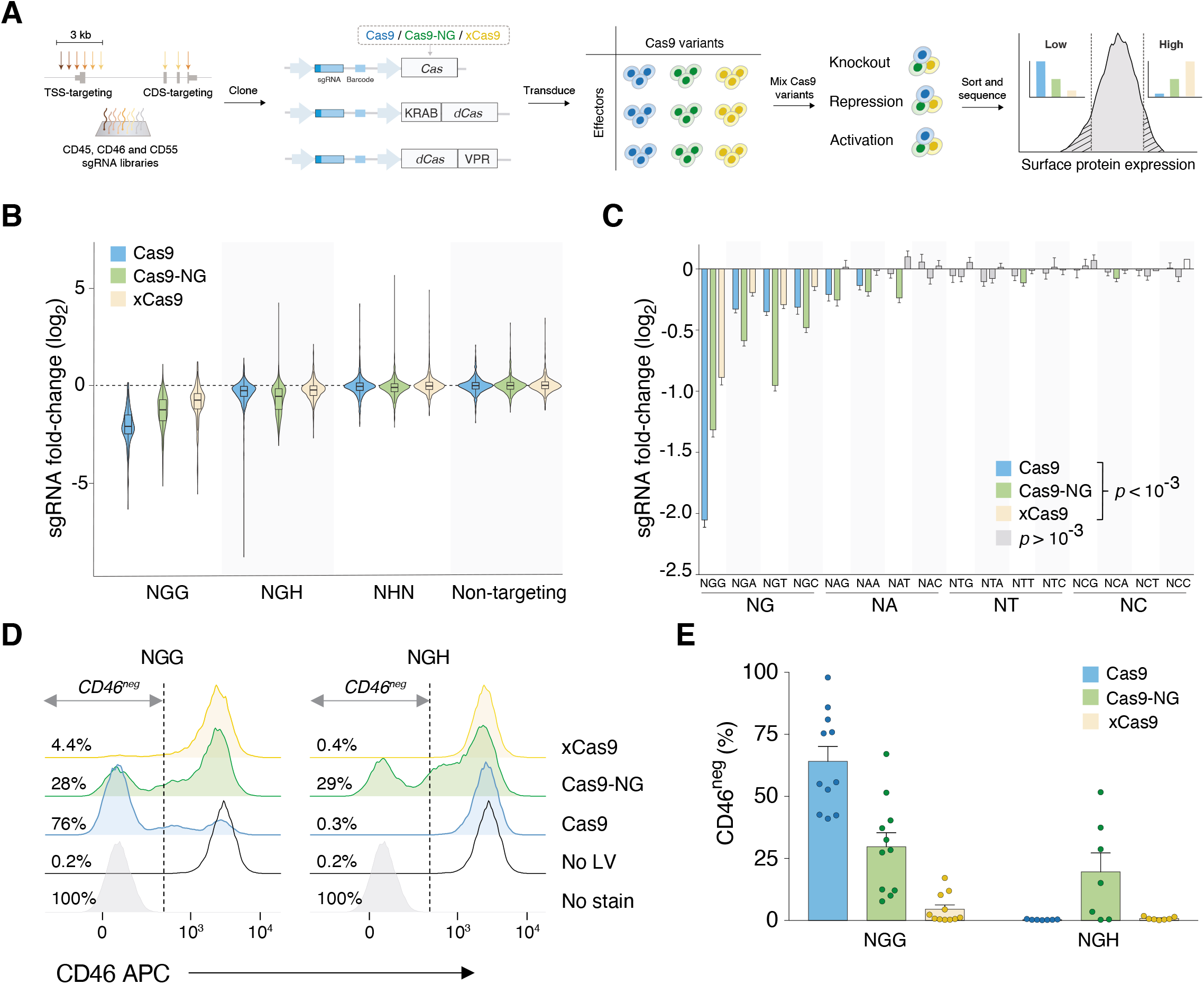
A high-throughput, pooled competition assay for PAM-flexible Cas9 variants. **A**) Gene-specific sgRNA libraries were cloned into lentiviral plasmids containing barcoded Cas9 effectors. After library transduction, K562 cells were sorted by target gene expression level into high- and low-expressing bins and the relative frequency of sgRNA-Cas9 barcode pairs in both bins was compared. **B**) Fold-change of sgRNA representation in cell populations expressing a high level of the target gene compared to low-expressing cells, combined for all three genes tested. Only sgRNAs targeting CDS exons are shown. **C**) Fold-change of sgRNA representation grouped by two- and three-nucleotide PAMs. Statistical significance was determined by comparing foldchange of sgRNAs associated with a particular PAM to a respective non-targeting control using two-tailed Student’s t-test with Bonferroni correction for multiple hypothesis testing. Error bars indicate standard error of the mean. Only statistically significant PAM/Cas9 combinations are shown in color. **D**) CD46^neg^ gate, indicated with a dashed line, was set based on K562 autofluorescence. Numbers displayed next to histograms indicate the percentage of cells in CD46^neg^ gate. LV – lentivirus. **E**) Quantification of CD46 knock-out in K562 cell line by lentiviral transduction of Cas9 nucleases and sgRNAs associated with NGG or NGH PAMs. Error bars indicate standard error of the mean.

To build a sufficiently large dataset, we selected single-guide RNAs (sgRNAs) at thousands of target sites spanning all possible three-nucleotide PAM combinations. Specifically, we designed three sgRNA libraries targeting the genes CD45, CD46 and CD55, which encode cell surface markers that can be detected by antibody labeling, and are expressed in human K562 cells (**Figure S1B,C**). For each gene-specific library, we selected sgRNAs that either target coding exons (CDS) or target within a 3 kb region flanking the transcription start site (TSS) (**Figure 1A**). Combining TSS and CDS targeting sgRNAs in a single library enabled us to use the same library to test for CRISPR nuclease activity (assaying gene disruption) and transcriptional modulation via CRISPRi or CRISPRa. In the target regions (CDS and TSS), we selected all available NGN PAMs and equal numbers of NHN PAMs (**Table S1**). In total, we synthesized 6,713 sgRNAs targeting these three genes. Each gene-specific library also included 250 sgRNAs that are predicted to not target anywhere in the human genome as negative controls (Sanjana et al., 2014).

The libraries were cloned into a lentiviral plasmid containing a Cas9 variant (WT, Cas9-NG or xCas9) and a six-nucleotide barcode specific for the particular Cas9 variant and given modality (nuclease, repression or activation). This plasmid design allowed us to determine simultaneously the sgRNA and Cas9 effector (barcode) identities by high-throughput Illumina sequencing (**Figure S1A**). Recently, several groups have reported lentiviral recombination between pseudodiploid viral RNAs as a function of distance within the viral RNA genome, which results in barcode swapping after transduction (Feldman et al., 2018; Hegde et al., 2018; Hill et al., 2018; Xie et al., 2018). To avoid these issues, we cloned and produced lentivirus separately for all 27 combinations of sgRNA libraries (CD45, CD46, CD55), Cas9 variants (WT, Cas9-NG or xCas9) and effector domains (nuclease, CRISPRi, CRISPRa). We separately transduced these libraries at a low multiplicity of infection into human K562 cells.

Following puromycin selection of transduced cells, we pooled together an equal number of cells transduced with different enzymes (WT, Cas9-NG or xCas9), performed antibody staining for each cell-surface protein, and sorted them by target expression via fluorescence-assisted cell sorting (FACS) (**Figure S1D**). Pooling the cells just prior to antibody labelling and sorting allowed us to compare the efficiency of each enzyme in a direct competition-like assay, as well as to tightly control the ratios of cells transduced with each enzyme in the pre-sort input, to ensure no prior bias towards any Cas9 variant (**Figure S3**). The relative frequency of every sgRNA-Cas9 variant pair from the top bin (highest expression) was then divided by its corresponding frequency in the bottom bin (lowest expression) to calculate the fold-change of sgRNAs associated with a particular PAM. In most cases, we found that the sgRNA distributions between Cas9 libraries in the mixed, pre-sort samples were tightly correlated (**Figure S4**).

### Cas9-NG targets NGH PAMs with 2-to 4-fold lower nuclease activity than Cas9 at NGG PAMs

We first performed the CRISPR competition screens using catalytically-active nucleases and compared the fold-change of sgRNAs targeting coding exons (*n* = 2,107 sgRNAs). Across all three cell-surface proteins, we observed the greatest fold-change for target sites with the canonical NGG PAM using the WT Cas9 enzyme (**Figure 1B; Figure S5A** shows each gene separately). Compared to WT Cas9, we found that the mean relative knock-out activity of Cas9-NG was 64% of WT and xCas9 was 43% of WT. For NGH PAMs, Cas9-NG provided the best overall knockout (**Figure 1B**). Unexpectedly, xCas9 was not significantly better than WT Cas9 at NGH PAMs. In contrast to CDS-targeting sgRNAs, sgRNAs targeting upstream noncoding regions for each of the three cell-surface proteins displayed only a minimal change in representation (**Figure S5B**).

To further dissect Cas9 variant activity at specific PAMs and to discover potentially targetable non-NG PAMs, we next examined all possible nucleotide combinations at PAM positions 2 and 3 (**Figure 1C**). While WT Cas9 showed the strongest activity at NGG PAMs, it was also capable of targeting endogenous genomic loci with all three NGH PAMs, albeit with greatly reduced activity. In addition to NGH PAMs, WT Cas9 showed significant recognition of NAG and NAA PAMs. Other groups have previously reported limited Cas9 nuclease activity in human cells at NAG PAMs, thus highlighting the sensitivity of our assay (Hsu et al., 2013; Zhang et al., 2015). Surprisingly, we found that xCas9 performed worse than WT Cas9 at all 3 NGH PAMs while PAM-flexible Cas9-NG was considerably more active than WT Cas9 or xCas9. Among NGH sites, Cas9-NG showed greatest activity at NGT PAMs and lowest activity at NGC PAMs, as reported previously (Nishimasu et al., 2018). In our screen, we also found that Cas9-NG was active at some non-NG PAMs, in particular at NAD (D = A, G or T) PAMs.

To further validate our pooled comparison, we targeted the CD46 gene in K562 cells with 18 individual sgRNAs at NGG and NGH PAMs using all 3 enzymes and quantified protein expression via FACS. To minimize bias due to sgRNA nucleotide composition, we designed sgRNAs targeting NGH PAMs to be shifted one nucleotide downstream from the corresponding NGG PAM-targeting sgRNAs. Following lentiviral transduction and selection, we measured the knockout efficiency by flow cytometry (**Figure 1D,E**). We observed robust gene knockout induced by WT Cas9 and sgRNAs targeting NGG PAMs with 64% of cells having a CD46^null^ phenotype. Cas9-NG at NGG PAMs induced full knockout at 46% efficiency of WT Cas9, followed by xCas9 at 7% of WT. At NGH PAMs, we could not detect any knockout above background induced by either WT Cas9 or xCas9; Cas9-NG activity at NGH PAMs was at 66% of its activity at corresponding NGG sites. Furthermore, xCas9 activity at NGG or NGH PAMs could not be rescued by increasing the editing time – even at day 21 post-transduction, knockout frequency with the best NGG sgRNA reached only 25% of knockout observed with Cas9-NG (**Figure S6**).

Interestingly, we noticed a difference in knockout kinetics between wild-type Cas9 and Cas9-NG. While knockout efficiency of Cas9-NG (at both NGG and NGH PAM sites) sharply increased between days 4 and 14 post-transduction, wild-type Cas9 activity reached levels close to saturation already at day 4. Finally, both Cas9-NG and xCas9 showed high variability in knockout efficiency between different sgRNAs, ranging from no detectable activity up to a maximum of 17% (xCas9) or 70% (Cas9-NG) CD46^neg^ cells. This observation highlights the advantage of our approach: testing thousands of sgRNAs in parallel can reduce target site-specific bias by averaging over many target sites.

We also measured the editing efficiency at the DNA level by high-throughput amplicon sequencing and we observed that the frequency of alleles with insertions or deletions (indels) correlated well with protein expression from flow cytometry (*r^2^* = 0.93, **Figure 2A,B**). Furthermore, there was no significant difference between the three Cas9 variants with regard to their preferences for insertions or deletions or to the mean indel size among edited alleles (**Figure 2C-E**).

**Figure 2.**
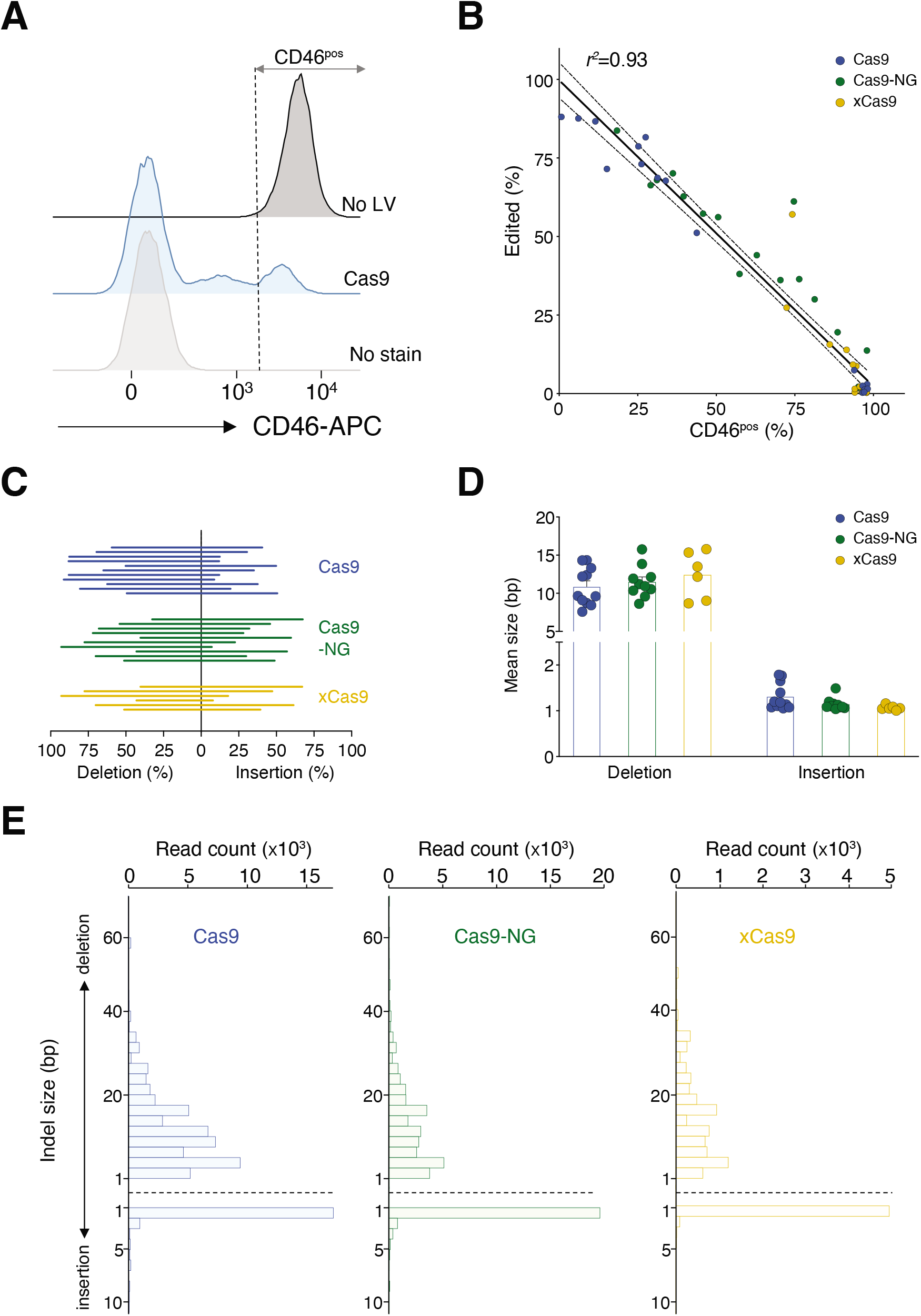
Characterization of indel mutations produced by active Cas9 variants. **A**) Gating strategy for enumeration of K562 cells expressing the wild-type level of CD46 protein. No LV – no lentivirus. **B**) Correlation between the frequency of alleles containing indels and the frequency of cells expressing the wild-type levels of CD46 protein. Dashed lines indicate 95% confidence intervals around the linear regression curve. NGG and NGH sgRNAs are included. *r*^2^ is the Pearson coefficient of determination. **C**) Relative frequency of deletions and insertions among edited alleles. Each line represents one sgRNA. Only NGG sgRNAs with >5% edited alleles are included. **D**) Mean deletion and insertion sizes per Cas9 variant. Each data point represents the mean indel size for one sgRNA. Error bars indicate SEM. Only NGG sgRNAs with >5% edited reads are included. **E**) Indel sizes (among edited reads) for each Cas9 variant.

### Cas9-NG, but not xCas9 or WT Cas9, efficiently modulates gene expression at NGH PAMs

CRISPR nuclease activity is a two-step process: first, the Cas9-sgRNA complex binds the target DNA and second, it undergoes a conformational change which enables double-strand break formation (Nishimasu et al., 2018; Wu et al., 2014). In contrast, CRISPR transcriptional modulation only requires Cas9 sgRNA binding in the target region to enable recruitment of transcriptional repressors or activators. We hypothesized that xCas9, which showed suboptimal performance as a nuclease, might perform better in context of CRISPRi and CRISPRa because it was evolved via selection for DNA binding without cleavage. In the phage-based evolution and selection assay used to derive xCas9, nuclease-null Cas9 (dCas9) was fused to an *E. coli* RNA polymerase and targeted upstream of an essential gene for phage replication (Hu et al., 2018). In that study, xCas9 was shown to have, on average, a 12-fold increase in activity in human cells over WT Cas9 when fused to the VPR transcriptional activator (Hu et al., 2018). Given our previous results with xCas9 nuclease, we wanted to determine if dCas9 variants of the 3 enzymes fused to transcriptional activators and repressors would result in greater activity at NGH PAMs.

For this purpose, we first examined sgRNAs for all NGG PAMs tiling the 3 kb region surrounding the gene’s primary TSS to identify the optimal target region for subsequent analysis and comparison across all PAMs. In general, we found that the optimal CRISPRi window was shifted downstream of the optimal CRISPRa window by ~120 bp, possibly resulting from the interference of the bound Cas9 complex with the assembly of transcriptional machinery at the TSS (**Figure 3A, Figure S7**). Previously, Doench and colleagues reported that for CRISPR inhibition, the optimal targeting window is between +25 and +75 bp downstream of the TSS while for CRISPRa the optimal window lies between −150 and −75 bp upstream of the TSS (Sanson et al., 2018). We found similar windows for optimal CRISPRi and CRISPRa transcriptional modulation with peak CRISPRi inhibition downstream (3’) of peak CRISPRa activation. In addition, our screen data showed multiple peaks that aligned with particular transcript isoforms, suggesting that sgRNA positioning could preferentially activate or repress transcription from a particular TSS.

**Figure 3.**
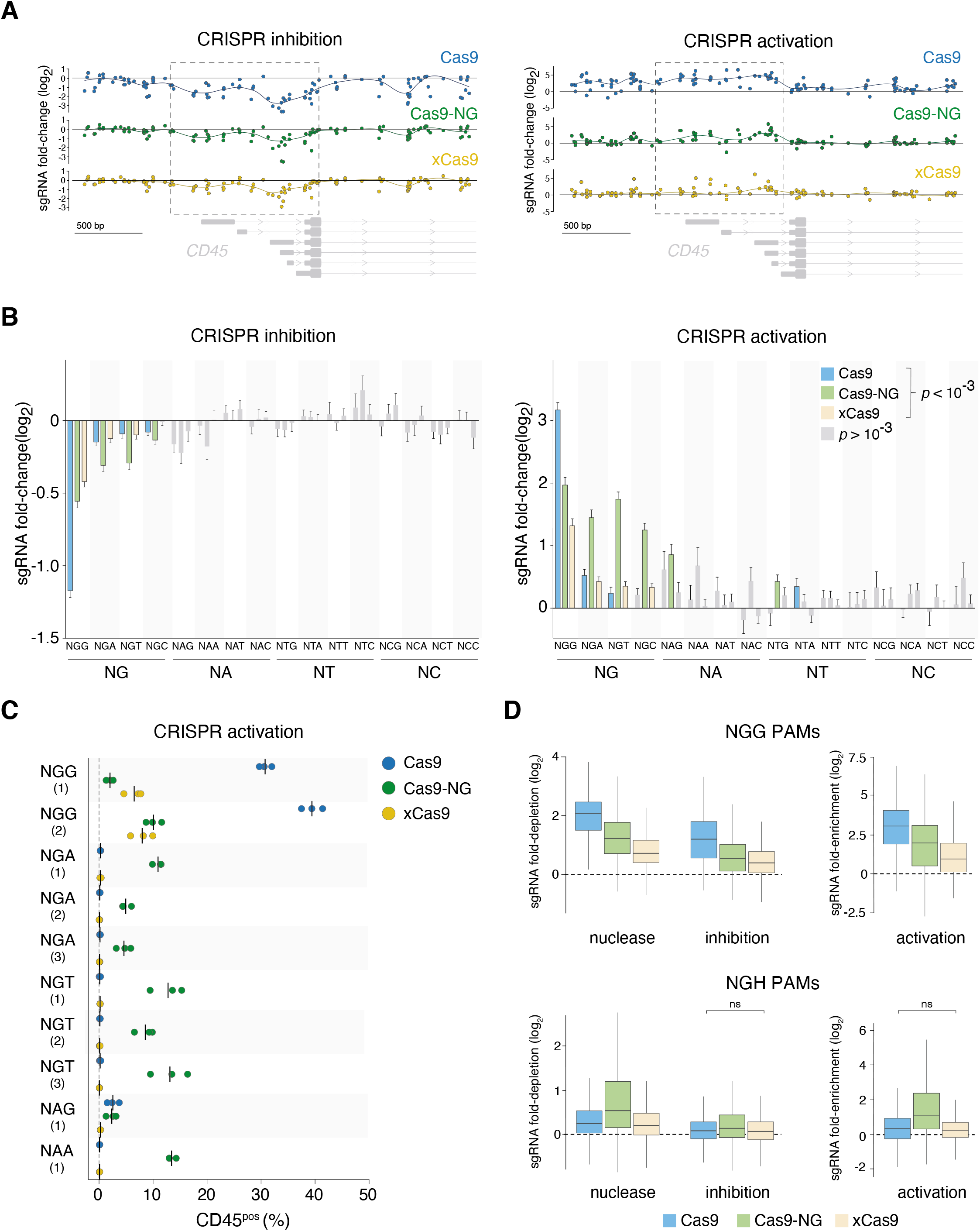
CRISPRi and CRISPRa transcriptional modulation using PAM-flexible Cas9 variants. **A)** Fold-change of sgRNAs targeting the 3 kb region surrounding the primary TSS of *CD45* gene. Only sgRNAs associated with an NGG PAM are displayed here (*n* = 123 sgRNAs). The regions with the strongest NGG sgRNA activity (indicated with dashed lines) were used to select sgRNAs (all PAMs) for subsequent analyses. *CD45* transcript isoforms (PTPRC-204, PTPRC-215, PRPRC-201, PTPRC-209, PTPRC-206, PTPRC-207) are shown in grey. **B**) Foldchange of sgRNA representation grouped by two- and three-nucleotide PAM categories. Statistical significance was determined by comparing fold-change of sgRNAs associated with a particular PAM to a respective non-targeting control using two-sided Student’s *t*-test with Bonferroni correction. Error bars indicate standard error of the mean. Only statistically significant PAM/Cas9 combinations are shown in color. CRISPRi: *n* = 2,165 sgRNAs; CRISPRa: *n* = 1,980 sgRNAs. **C**) CD45 expression following CRISPR activation in the CD45^neg^ human A375 cell line. Only sgRNAs resulting in >1% CD45^pos^ cells with at least one Cas9 variant are displayed (see **Figure S8** for data from all sgRNAs tested). Mean and individual values from three independent experiments are shown. For NAG and NAA PAMs, only one out of three sgRNAs tested resulted in >1% CD45^pos^ cells. **D**) Comparison of wild-type Cas9 and two PAM-flexible Cas9 variants across all three modalities tested in high-throughput screens. Fold-enrichment was calculated based on the sgRNA frequency in the top bin over bottom bin; fold-depletion was calculated based on the sgRNA frequency in the bottom bin over top bin. Only non-significant comparisons (ns, *p*>0.05) are indicated; all other differences (between enzymes, within modalities) are significant.

Overall, we observed that WT dCas9 produced the strongest effect on transcriptional modulation at NGG PAMs (**Figure 3B, Figure S8**). At NGH PAMs, dCas9-NG outperformed the other enzymes while dxCas9 had a similarly low activity to WT dCas9 at these PAMs, suggesting that xCas9 may not bind NGH PAMs as strongly as Cas9-NG. We also detected significant activity of dCas9-NG at unconventional NAD PAMs in context of CRISPR activation. This result is in agreement with our previous finding of Cas9-NG nuclease activity at NAD PAMs (**Figure 1C**). As expected, there was no apparent difference between PAM sites or Cas9 variants when we looked at the fold-change of sgRNAs targeting CDS exons distant from the TSS (**Figure S9**).

To further validate the pooled competition screen results, we targeted CD45 gene expression using 23 individual sgRNAs in two CD45^neg^ cell lines, A375 (**Figure 3C, Figure S10**) and HEK293T (**Figure S11**), using CRISPRa. In addition to NGN PAMs, we also used unconventional NAD PAMs identified from our CRISPRn and CRISPRa screen analyses. WT Cas9 outperformed the PAM-flexible enzymes at the two NGG sites tested. For NGH PAMs, Cas9-NG demonstrated greater activity at NGT over NGA PAMs, in agreement with the pooled screen. We also detected Cas9-NG activity at one out of three NAG and one out of three NAA sites tested. Although xCas9 showed similar activity at NGG sites to Cas9-NG, there was no detectable CRISPRa-driven CD45 protein expression when targeting non-NGG sites with xCas9.

We next computed the relative activity of all three Cas9 enzymes at NGG and NGH PAMs, across all three modalities tested (nuclease, transcriptional activation, transcriptional repression), integrating data from nine separate CRISPR competition screens (**Figure 3D**). At NGG PAMs, the strongest effector was WT Cas9 regardless of the modality, followed by Cas9-NG and then xCas9. At NGH PAMs, Cas9-NG showed significantly stronger activity than either WT Cas9 or xCas9. We found that xCas9 activity was not statistically different from WT Cas9 for transcriptional activation and repression at NGH PAMs; for nuclease activity, xCas9 was slightly, but significantly, weaker than WT Cas9. Overall, in three cell lines tested, Cas9-NG significantly outperformed xCas9 at NGH sites (**Figure S12**). Similar results were obtained using both lentiviral transduction and plasmid transfection (data not shown).

### Introduction of Cas9-NG mutations in xCas9 partially rescues nuclease activity and increases transcriptional activation at NGH PAMs

Our high-throughput CRISPR pooled competition screens and arrayed sgRNA validation data indicated that Cas9-NG is active for all modalities at NGN PAMs, albeit to a lesser extent than WT Cas9 at NGG sites. We also found that xCas9 had the poorest performance at virtually all PAMs and for all modalities. Due to this marked difference in Cas9-NG and xCas9 activity, we examined the position of the mutations in both Cas9 variants (**Figure 4A**). The mutations in Cas9-NG cluster together in the PAM-interacting domain, as expected from structure-guided design. Conversely, xCas9 mutations, generated through directed evolution, are spread throughout the protein, with only one mutated residue (E1219) in common with Cas9-NG. Given their disparate positions in the protein, we wondered if it might be possible to rescue xCas9 activity using mutations from Cas9-NG. For this purpose, we created a new Cas9 variant that combines mutations from both xCas9 and Cas9-NG (with E1219F from Cas9-NG) and termed this novel variant xCas9-NG.

**Figure 4.**
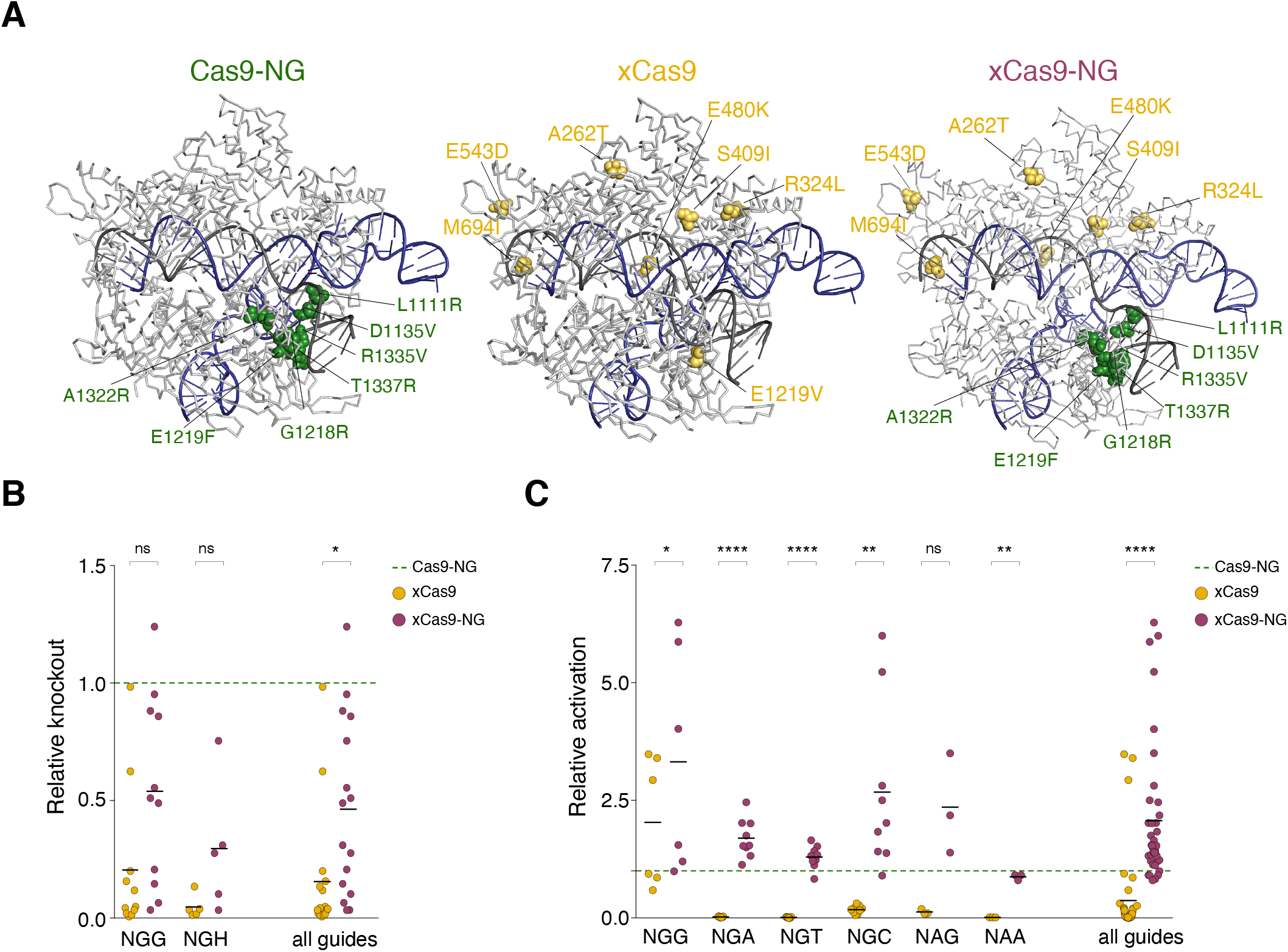
Cas9-NG mutations partially rescue xCas9 nuclease activity and results in an improved PAM-flexible Cas9 enzyme for transcriptional activation. **A**) Crystal structures of Cas9 mutants. xCas9 mutations are shown on a wild-type Cas9 structure (PDB ID: 4un3, (Anders et al., 2014). xCas9-NG mutations are displayed on a Cas9-NG structure (PDB ID: 6ai6,(Nishimasu et al., 2018). The sgRNA is shown in black and the target DNA is shown in blue. **B**) Knockout and **C**) activation activity for individual sgRNAs with target sites with the indicated PAMs. Each xCas9 or xCas9-NG experiment was normalized to Cas9-NG on a per-sgRNA basis. Only sgRNAs resulting in >1% knock-out or activation are shown. Non-normalized data for xCas9 and Cas9-NG were displayed in **Figures 1E** and **2C** and are included here for comparison with xCas9-NG. ns, *p* > 0.05; \**p* < 0.05, \*\**p* < 0.01, \*\*\*\**p* < 0.0001.

Using Cas9-NG as a baseline, we compared xCas9 and xCas9-NG nucleases using several sgRNAs to target CD46 in K562 cells at both NGG and NGH PAMs (**Figure 4B**). For comparison, we normalized the effects of each sgRNA with either xCas9 or xCas9-NG to the same sgRNA with Cas9-NG. The ability of xCas9-NG to drive gene knockout was overall three times stronger than that of xCas9 but remained at ~50% of Cas9-NG activity. For CRISPRa, the mean dxCas9-NG activation was on average two-fold greater than dCas9-NG and over five-fold greater than xCas9, across virtually all sgRNAs and for all NGN PAMs (**Figure 4C**). In particular, dxCas9-NG had 2.7-fold higher activation than dCas9-NG at NGC PAMs, which is especially important given that Cas9-NG had very low activity at these PAMs. In an independent cell line (HEK293FT), we confirmed that xCas9-NG resulted in significantly greater transcriptional activation, albeit to a lesser extent than in A375 cell line, than either existing PAM-flexible Cas9 variant (**Figure S13**). For CRISPR inhibition, xCas9-NG outperformed xCas9 with virtually every sgRNA tested, as well as outperformed Cas9-NG with one out of two NGG sgRNAs (**Figure S14A**). Overall, when looking at all 6 NGN sites tested, xCas9-NG drove the same level of transcriptional repression as Cas9-NG (**Figure S14B**). Thus, xCas9-NG appears to be a generally stronger transcriptional activator and an equal transcriptional repressor as Cas9-NG which may be possibly due to mechanistic differences between CRISPR activation and repression. Further kinetic and biochemical studies are warranted to fully elucidate the mechanistic features of transcriptional modulation and binding at specific NGH PAMs.

## Discussion

Taken together, we performed nine independent CRISPR competition screens, spanning three endogenously expressed human genes and three CRISPR modalities, to assess the efficacy of recently-described PAM-flexible Cas9 variants at different PAM sites. These are the first pooled CRISPR screens using xCas9 or Cas9-NG, testing thousands of endogenous genomic loci in a massively-parallel manner. By combining cells transduced with all three Cas9 variants prior to FACS, we were able to perform a pooled comparison where each variant competes against other variants. This high-throughput CRISPR competition screen provides a general method of assessing relative efficacies of PAM-flexible Cas9 variants and provides a far richer dataset than previous work with only a few target sites (Hu et al., 2018; Nishimasu et al., 2018). While this screen was not designed to discover sequence features determining the on-target efficiency of PAM-flexible Cas9 enzymes, that could be achieved by scaling up the number of assayed sgRNAs.

We showed that the mutations that increase PAM flexibility of Cas9 lead to decreased activity of these enzymes at NGG target sites. This observation applies to both catalytically active and inactive Cas9 variants. When comparing Cas9 variants at target sites with NGH PAMs, we were surprised to discover that while Cas9-NG maintains a similar level of activity as for target sites with NGG PAMs, the activity of xCas9 was profoundly diminished. In fact, at target sites with NGH PAMs, xCas9 did not perform better than wild-type Cas9 across all modalities tested (nuclease, activation, and inhibition). The discrepancies between the results reported in this study and in the original xCas9 publication could potentially stem from differences in accessibility of the target sites, thus highlighting the need to test endogenous loci for meaningful comparisons. Recent studies in plants (Ge et al., 2019; Hua et al., 2019; Negishi et al., 2019; Wang et al., 2019; Zhong et al., 2019) have shown that the overall efficiency of indel formation and base editing at non-NGG sites is much higher for Cas9-NG than for xCas9, supporting our findings in the mammalian context. Furthermore, David Liu and colleagues recently demonstrated that Cas9-NG base editors outperform xCas9 base editors at target sites with NGH PAMs and observed very low or no editing at the vast majority of loci tested when using xCas9 (Huang et al., 2019).

Structural studies have shown that the mechanisms behind relaxed PAM recognition by xCas9 and Cas9-NG are considerably different. In case of Cas9-NG, NG PAM recognition is enabled by mutating both the R1335 residue interacting with the third nucleobase of the PAM (dG3), and E1219, which stabilizes R1335. The remaining five mutations are introduced to enhance Cas9-NG binding to the now smaller, two-nucleobase PAM (Nishimasu et al., 2018). Conversely, in xCas9 the R1335-dG3 interaction is disrupted indirectly, by abrogating the E1219-R1335 interaction and allowing R1335 to adopt multiple conformations (Guo et al., 2019). The remaining xCas9 mutations are located in the recognition (REC) lobes and result in the conformational change of Cas9 binding to DNA.

Given these differences, we investigated how the change of REC lobes conformation (xCas9 mutations) would affect the editing activity of the enzyme when combined with enhanced binding to the two-nucleobase PAM (Cas9-NG mutations). This new Cas9 variant, termed xCas9-NG, showed improved nuclease activity compared to xCas9, presumably due to stronger interactions with the PAM, although it did not fully rescue nuclease activity to the Cas9-NG level. In contrast, we also found that xCas9-NG was superior to both xCas9 and Cas9-NG for transcriptional modulation, possibly indicating that a more relaxed REC lobe interaction with target DNA allows for easier access of the recruited transcriptional machinery. Over the entire human exome and functional non-coding regions, the relaxed PAM constraints of xCas9-NG enable a significantly larger target space (**Figure S15A-D**), especially when considering the additional NAD PAMs found in our screens.

As none of the three PAM flexible Cas9 mutants were capable of matching the efficacy of wildtype Cas9 at NGG PAM sites, relaxing PAM interactions through these mutations likely incurs a fitness cost in enzyme performance. New strategies are needed for designing efficient, PAM-flexible (or perhaps even PAM-independent) Cas9 enzymes. The CRISPR competition screen presented here provides a robust and scalable platform for future benchmarking of different genome editing enzymes prior to their implementation in research, clinical or industrial applications.

## Methods and materials

### Cell culture

K562 and A375 cell lines were obtained from ATCC. HEK 293FT cells were obtained from Thermo Scientific. K562 cells were cultured in Iscove’s Modified Dulbecco’s Medium (IMDM); A375 and HEK293FT cells were cultured in Dulbecco’s Modified Eagle Medium (DMEM). All media were from Caisson Labs. Media were supplemented with 10% Serum Plus II Medium Supplement (Sigma-Aldrich). Cells were regularly passaged and tested for presence of mycoplasma contamination (MycoAlert Plus Mycoplasma Detection Kit, Lonza).

### Plasmid design

In order to enable a meaningful comparison between different Cas9 variants, we used the human codon optimized Cas9 from lentiCRISPR v2 plasmid (Addgene 52961, Sanjana et al., 2014) as background for xCas9 and Cas9-NG mutations. xCas9 (also known as xCas3.7) mutations are as follows: A262T, R324L, S409I, E480K, E543D, M694I and E1219V (Hu et al., 2018). Cas9-NG mutations are: L1111R, D1135V, G1218R, E1219F, A1322R, R1335V, T1337R (Nishimasu et al., 2018). xCas9-NG mutations are: A262T, R324L, S409I, E480K, E543D, M694I, L1111R, D1135V, G1218R, E1219F, A1322R, R1335V and T1337R. For transcriptional modulation, Cas9 variants contained additional D10A and H840A mutations to make them catalytically inactive. KRAB domain was derived from pHAGE EF1α dCas9-KRAB (Addgene 50919, Kearns et al., 2014). VPR complex was derived from lenti-EF1a-dCas9-VPR-Puro (Addgene 99373, Ho et al., 2017) and modified to abolish BsmBI restriction sites. sgRNA scaffold was modified to improve its stability and Cas9 binding (F+E modification, Chen et al., 2013). Finally, we inserted a six-nucleotide barcode between the sgRNA scaffold and EFS promoter to act as an identifier for Cas9 variant and CRISPR modality (**Figure S1A**). All cloning was performed by Gibson Assembly using recombinase-deficient NEB Stable cells (all from New England Biolabs). Cloned inserts were fully validated by Sanger sequencing (Eton Bioscience). All plasmids have been deposited on Addgene.

### Lentiviral sgRNA library design and cloning

The sgRNAs targeting the 3 kb region surrounding the TSS and constitutive protein-coding exons were chosen to include all possible 20-mer sequences upstream of an NG PAM sequence, and equal numbers of 20-mer sequences upstream of NH PAM sequences. Primary TSS and exon annotations were obtained from the UCSC Genome Browser based on the hg38 genome assembly. We also included 250 non-targeting sgRNAs from the GeCKO v2 library (Sanjana et al., 2014) as a negative control in each library. **Table S1** specifies the number of sgRNAs per category. The sgRNA library was synthesized as an oligo pool of 103 nt oligos (Twist Bioscience) and cloned using Gibson Assembly (New England Biolabs) into Esp3I-digested (ThermoScientific/Fermentas) lentiviral transfer plasmids containing Cas9 effectors. The cloned libraries were individually amplified by electroporation into Endura ElectroCompetent cells (Lucigen). Using dilution plates for colony counting, we verified that all libraries were cloned with ≥1,000 × library coverage. Plasmids with cloned libraries were sequenced to confirm representation (MiSeq).

### Production of lentivirus and transduction

Lentivirus was produced by polyethylenimine linear MW 25000 (Polysciences 23966) transfection of HEK 293FT cells with the transfer plasmid containing a barcoded Cas9 effector and sgRNA library, packaging plasmid psPAX2 (Addgene 12260) and envelope plasmid pMD2.G (Addgene 12259). After 72 h post-transfection, cell media containing lentiviral particles was harvested and filtered through 0.45 μm filter Steriflip-HV (Millipore SE1M003M00). Each sgRNA library and Cas9 effector combination was transduced into K562 cells individually, to avoid barcode swapping, and thus Cas9 misidentification, during lentiviral integration (Xie et al., 2018). In total we produced 27 individual lentiviral libraries and transduced them into separately into K562 cells. The transduction was performed at a multiplicity of infection (MOI) ~ 0.4 to minimize the fraction of cells with multiple sgRNAs. We maintained 1,000× coverage of each sgRNA library. Transduced cells were selected with 1 μg ml^-1^ puromycin for at least 7 days after transduction. During the course of the screen the cells were maintained at numbers ensuring >1,000× representation of the library. Transduced cells were maintained as 27 separate cell cultures for 14 days. At day 14 post-transduction, cells transduced with the sgRNA library targeting the same gene and the same CRISPR modality (but different Cas9 variants) were combined in equal numbers, resulting in 9 separate cell pools for screening, and then analyzed and sorted via FACS. All cell counting was done using a Cellometer Auto T4 counter (Nexcelom).

For arrayed CD46 knockout validation in K562 and A375 cells, sgRNAs targeting exons 2 and 3 of *CD46* gene were designed in benchling software as 20-mers upstream of an NGG PAM, or by shifting +1 bp upstream, as 20-mers upstream of an NGH PAM (**Table S2**). The individual sgRNAs were cloned into lentiviral transfer plasmids encoding Cas9 variants and transduced into K562/A375 cells at MOI ~ 0.5. K562 cells were assessed for CD46 knockout by flow cytometry on day 14 after transduction. At this timepoint an aliquot of cells was also collected for genomic DNA (gDNA) extraction. A375 cells were assessed for CD46 knockout by flow cytometry on days 4, 7, 14 and 21 after transduction.

For arrayed CRISPR inhibition validation, we selected guide RNAs from sequences included in the screen library (NG PAMs) or designed to target within close proximity to NG PAM sgRNAs (NH PAM). The sequences of sgRNAs are listed in **Table S2.** The individual sgRNAs were cloned into lentiviral transfer plasmids encoding Cas9 variants and transduced into K562 cells at MOI ~ 0.5. K562 cells were assessed for CD45 knockdown by flow cytometry on day 14 after transduction.

### Transfection

For arrayed CRISPR activation validation, sgRNA-specifying oligos were either obtained from sgRNA sequences included in the screen library (NG PAMs) or designed to target within close proximity to NG PAM sgRNAs (NH PAM). The sequences of sgRNAs are listed in **Table S2.** The individual sgRNAs were cloned into a sgRNA-only plasmid with the F+E scaffold modification (Chen et al., 2013) and co-transfected with plasmids containing Cas9 effectors into A375 or HEK 293FT cells using Lipofectamine 2000 (ThermoFisher 11668019). The transfected cells were selected with 2 μg ml^-1^ puromycin for 72 h. At day 4 post-transfection, the cells were assessed for CD45 expression by flow cytometry.

### Protein expression

HEK293FT cells were transiently transfected with equal amounts of Cas9 variants expression vectors. At 24 hours post-transfection, the cells were collected, lysed with TNE buffer (10 mM Tris-HCl, pH 7.4, 150 mM NaCl, 1mM EDTA, 1% Nonidet P-40) supplemented with protease inhibitor cocktail (Bimake B14001) for 1 hour on ice. Cells lysates were spun for 10 min at 10,000 g, and supernatants were used to determine the protein concentration for each sample using the BCA assay (ThermoFisher 23227). Equal amounts of whole cell lysates (20 μg protein per sample) were denatured in Tris-Glycine SDS Sample buffer (ThermoFisher LC2676), and loaded on a Novex 4-20% Tris-Glycine gel (ThermoFisher XP04205BOX). PageRuler pre-stained protein ladder (ThermoFisher 26616) was used to determine the protein size. The gel was run in 1x Tris-Glycine-SDS buffer (IBI Scientific IBI01160) for 20 min at 80V, and then for additional 100 min at 120V. Proteins were transferred on a nitrocellulose membrane (BioRad 1620112) in presence of prechilled 1x Tris-Glycine transfer buffer (FisherSci LC3675) supplemented with 20% methanol for 100 min at 100V. Immunoblots were blocked with 5% skim milk dissolved in 1x PBS + 1% Tween 20 (PBST), washed well with PBST and incubated overnight at 4°C separately with the following primary antibodies: mouse anti-2A peptide, clone 3H4 (1 μg/mL, Millipore MABS2005); rabbit anti-GAPDH 14C10 (0.1 μg/mL, Cell Signaling 2118S). Following the primary antibody, the blots were incubated with IRDye 680RD donkey anti-rabbit (0.2 μg/mL, LI-COR 926-68073) or with IRDye 800CW donkey anti-mouse (0.2 μg/mL, LI-COR 926-32212). The blots were imaged using Odyssey CLx (LI-COR). Band intensity quantification was performed using ImageJ version 1.51.

### Flow cytometry and FACS

For CRISPR library sorting, >10^8^ cells were taken for antibody staining (~10,000× library representation). We set aside 10^7^ cells for the pre-sort control (~1,000× coverage). After harvesting the cells and removing leftover medium by washing with PBS, the cells were stained for 5 minutes at room temperature with LIVE/DEAD Fixable Violet Dead Stain Kit (ThermoFisher L34864). Subsequently, the cells were stained with antibodies for 20 minutes on ice. The following antibodies were used: CD45-PE (clone 2D1), CD46-APC (clone TRA-2-10) or CD55-APC (clone JS11). All antibodies were purchased pre-conjugated from BioLegend. Cells were washed with PBS to remove unbound antibodies prior to sorting. Cell acquisition and sorting was performed using a Sony SH800S cell sorter.

Sequential gating was performed as follows: 1) exclusion of debris based on forward and side scatter cell parameters, 2) doublet exclusion, and 3) dead cell exclusion (**Figure S1B**). The sorting gates were set based on the expression level of the target protein in sgRNA library-transduced cells (top and bottom 15% of expression, **Figure S1D**). Typically, we achieved >500× library coverage within each sorted population.

### Pooled CRISPR competition assay sequencing

The sgRNA library preparation was performed as described before (Shalem et al., 2014). Briefly, gDNA was extracted using GeneJET DNA Purification Kit (Thermo Fisher Scientific). All of the extracted gDNA was then used in the first PCR reaction, in multiple reactions not exceeding 10 μg gDNA per 100 uL PCR reaction. Samples were then subjected to a second PCR to add sequencing adaptors and to barcode the samples. All PCR primers are listed in **Table S3**. PCR products were run on a 2% agarose gel and the correct size band was extracted. PCR products from different samples were then pooled together in equimolar ratios. Sequencing was performed on the NextSeq 500 instrument using the MidOutput Mode v2 with 75 bp paired-end reads (Illumina).

### Pooled CRISPR competition assay data analysis

The sgRNA sequences present in the sorted samples (read 1) as well as their corresponding barcodes indicating the Cas9 variant and CRISPR modality (read 2) were enumerated. sgRNA sequences were mapped to the reference sgRNA library with one mismatch allowed (bowtie −v 1 −m 1). Read numbers were normalized to the total number of reads per sample (with a pseudocount added to all sgRNAs) and log2-transformed. The median of non-targeting sgRNAs was calculated for each of the three Cas9 variants present in a sample. The median of non-targeting (NT) sgRNAs associated with each Cas9 was then used to normalize the sgRNA read counts associated with that Cas9. Finally, the fold-change of each NT-normalized sgRNA-Cas9 pair in top 15% bin was calculated over the NT-normalized sgRNA-Cas9 pair in the bottom 15% bin. Statistical significance was determined by two-sided Student’s *t*-test with Bonferroni correction (RStudio). For CRISPRi and CRISPRa screens, we needed to determine optimal windows around the TSS to pick the sgRNAs for subsequent analyses (i.e. to compare Cas9 variants across NGN PAMs, and to identify new functional NHN PAMs). Windows were selected to capture the peak region identified from the LOESS fit for all three enzymes, using only the NGG sgRNAs for strongest signal. The following parameters were chosen for LOESS fitting using the Gviz package (Hahne and Ivanek, 2016) in RStudio: span=0.2, evaluation=500, degree=10.

### Nuclease indel sequencing

For validation of arrayed CD46 knockout, genomic DNA was isolated using QuickExtract DNA Extraction Solution (Epicentre). Two sets of PCR primers were designed: first set was flanking the exons to be amplified and contained handles for the second PCR. The primers for the second PCR were handle-specific, and added Illumina sequencing adaptors and indexes (**Table S3**). PCR products of the correct size were extracted following agarose gel electrophoresis, combined in equimolar amounts and sequenced on the NextSeq 500 instrument using the MidOutput Mode v2 with 150 bp single-end reads (Illumina).

### Data analysis

Illumina single-end reads for CD46 genomic amplicons were analyzed using CRISPResso2 software (Clement et al., 2019) to quantify the fraction of reads containing editing at expected sites, and to determine the editing outcome in terms of indel type and size. Flow cytometry data was analyzed using FlowJo software. Visualization of Cas9 protein structures was performed in PyMOL software (PDB IDs: 4un3; 6ai6). All other data analysis was performed in GraphPad Prism 8 and RStudio. All correlation coefficients (*r*) and coefficients of determination (*r^2^*) are Pearson’s correlation. DNase I hypersensitivity (HS) sites in the K562 cell line were downloaded from ENCODE DNase Uniformly Processed Peaks from UCSC based on hg19 genome build.

### Data representation

In all boxplots, boxes indicate the median and interquartile ranges, with whiskers indicating either 1.5 times the interquartile range, or the most extreme data point outside the 1.5-fold interquartile. All transfection experiments show the mean of three replicate experiments, with error bars representing the standard error of mean.

### Data availability statement

Screen data are being deposited to GEO with an accession number GSE143892. All plasmids have been deposited on Addgene. Other data and materials that support the findings of this research are available from the corresponding author upon reasonable request.

## Acknowledgements

We thank the entire Sanjana laboratory for support and advice. We also thank the New York Genome Center for flow cytometry resources and the NYU Biology GenCore for sequencing resources. N.E.S. is supported by NYU and NYGC startup funds, NIH/NHGRI (R00HG008171, DP2HG010099), NIH/NCI (R01CA218668), DARPA (D18AP00053), the Sidney Kimmel Foundation, and the Brain and Behavior Foundation. M.L. is supported by the Hope Funds for Cancer Research postdoctoral fellowship.

## Author contributions

N.E.S. and M.L. conceived the project. N.E.S., M.L. and Z.D. designed the experiments. M.L., Z.D., X.X. and D.M. performed the experiments. M.L. and Z.D. analyzed the experiments. H.H.W. and X.G. assisted with screen data analysis and presentation. N.E.S. supervised the work. M.L., Z.D. and N.E.S. wrote the manuscript with input from all the authors.

## References

Anders, C., Niewoehner, O., Duerst, A., and Jinek, M. (2014). Structural basis of PAM-dependent target DNA recognition by the Cas9 endonuclease. Nature 513, 569–573.

Chavez, A., Tuttle, M., Pruitt, B.W., Ewen-Campen, B., Chari, R., Ter-Ovanesyan, D., Haque, S.J., Cecchi, R.J., Kowal, E.J.K., Buchthal, J., et al. (2016). Comparison of Cas9 activators in multiple species. Nature Methods 13, 563–567.

Chen, B., Gilbert, L.A., Cimini, B.A., Schnitzbauer, J., Zhang, W., Li, G.-W., Park, J., Blackburn, E.H., Weissman, J.S., Qi, L.S., et al. (2013). Dynamic Imaging of Genomic Loci in Living Human Cells by an Optimized CRISPR/Cas System. Cell 155, 1479–1491.

Clement, K., Rees, H., Canver, M.C., Gehrke, J.M., Farouni, R., Hsu, J.Y., Cole, M.A., Liu, D.R., Joung, J.K., Bauer, D. E., et al. (2019). CRISPResso2 provides accurate and rapid genome editing sequence analysis. Nature Biotechnology 37, 224–226.

Feldman, D., Singh, A., Garrity, A.J., and Blainey, P.C. (2018). Lentiviral co-packaging mitigates the effects of intermolecular recombination and multiple integrations in pooled genetic screens. BioRxiv.

Findlay, G.M., Boyle, E.A., Hause, R.J., Klein, J.C., and Shendure, J. (2014). Saturation editing of genomic regions by multiplex homology-directed repair. Nature 513, 120–123.

Ge, Z., Zheng, L., Zhao, Y., Jiang, J., Zhang, E.J., Liu, T., Gu, H., and Qu, L. (2019). Engineered xCas9 and SpCas9-NG variants broaden PAM recognition sites to generate mutations in *Arabidopsis* plants. Plant Biotechnology Journal.

Guo, M., Ren, K., Zhu, Y., Tang, Z., Wang, Y., Zhang, B., and Huang, Z. (2019). Structural insights into a high fidelity variant of SpCas9. Cell Research 29, 183–192.

Hahne, F., and Ivanek, R. (2016). Visualizing Genomic Data Using Gviz and Bioconductor. In Statistical Genomics, E. Mathé, and S. Davis, eds. (New York, NY: Springer New York), pp. 335–351.

Hegde, M., Strand, C., Hanna, R.E., and Doench, J.G. (2018). Uncoupling of sgRNAs from their associated barcodes during PCR amplification of combinatorial CRISPR screens. PLoS ONE 13, e0197547.

Hill, A.J., McFaline-Figueroa, J.L., Starita, L.M., Gasperini, M.J., Matreyek, K.A., Packer, J., Jackson, D., Shendure, J., and Trapnell, C. (2018). On the design of CRISPR-based single-cell molecular screens. Nature Methods 15, 271–274.

Ho, S.-M., Hartley, B.J., Flaherty, E., Rajarajan, P., Abdelaal, R., Obiorah, I., Barretto, N., Muhammad, H., Phatnani, H.P., Akbarian, S., et al. (2017). Evaluating Synthetic Activation and Repression of Neuropsychiatric-Related Genes in hiPSC-Derived NPCs, Neurons, and Astrocytes. Stem Cell Reports 9, 615–628.

Hsu, P.D., Scott, D.A., Weinstein, J.A., Ran, F.A., Konermann, S., Agarwala, V., Li, Y., Fine, E.J., Wu, X., Shalem, O., et al. (2013). DNA targeting specificity of RNA-guided Cas9 nucleases. Nature Biotechnology 31, 827–832.

Hu, J.H., Miller, S.M., Geurts, M.H., Tang, W., Chen, L., Sun, N., Zeina, C.M., Gao, X., Rees, H.A., Lin, Z., et al. (2018). Evolved Cas9 variants with broad PAM compatibility and high DNA specificity. Nature 556, 57–63.

Hua, K., Tao, X., Han, P., Wang, R., and Zhu, J.-K. (2019). Genome Engineering in Rice Using Cas9 Variants that Recognize NG PAM Sequences. Molecular Plant 12, 1003–1014.

Huang, T.P., Zhao, K.T., Miller, S.M., Gaudelli, N.M., Oakes, B.L., Fellmann, C., Savage, D.F., and Liu, D.R. (2019). Circularly permuted and PAM-modified Cas9 variants broaden the targeting scope of base editors. Nature Biotechnology 37, 626–631.

Jinek, M., Chylinski, K., Fonfara, I., Hauer, M., Doudna, J.A., and Charpentier, E. (2012). A Programmable Dual-RNA-Guided DNA Endonuclease in Adaptive Bacterial Immunity. Science 337, 816–821.

Kearns, N.A., Genga, R.M.J., Enuameh, M.S., Garber, M., Wolfe, S.A., and Maehr, R. (2014). Cas9 effector-mediated regulation of transcription and differentiation in human pluripotent stem cells. Development 141, 219–223.

Kim, D., Kim, J., Hur, J.K., Been, K.W., Yoon, S., and Kim, J.-S. (2016). Genome-wide analysis reveals specificities of Cpf1 endonucleases in human cells. Nature Biotechnology 34, 863–868.

Kleinstiver, B.P., Prew, M.S., Tsai, S.Q., Topkar, V.V., Nguyen, N.T., Zheng, Z., Gonzales, A.P.W., Li, Z., Peterson, R.T., Yeh, J.-R.J., et al. (2015). Engineered CRISPR-Cas9 nucleases with altered PAM specificities. Nature 523, 481–485.

Komor, A.C., Kim, Y.B., Packer, M.S., Zuris, J.A., and Liu, D.R. (2016). Programmable editing of a target base in genomic DNA without double-stranded DNA cleavage. Nature 533, 420–424.

Meier, J.A., Zhang, F., and Sanjana, N.E. (2017). GUIDES: sgRNA design for loss-of-function screens. Nature Methods 14, 831–832.

Negishi, K., Kaya, H., Abe, K., Hara, N., Saika, H., and Toki, S. (2019). An adenine base editor with expanded targeting scope using SpCas9-NG v1 in rice. Plant Biotechnology Journal.

Nishimasu, H., Shi, X., Ishiguro, S., Gao, L., Hirano, S., Okazaki, S., Noda, T., Abudayyeh, O.O., Gootenberg, J.S., Mori, H., et al. (2018). Engineered CRISPR-Cas9 nuclease with expanded targeting space. Science 361, 1259–1262.

Ran, F.A., Cong, L., Yan, W.X., Scott, D.A., Gootenberg, J.S., Kriz, A.J., Zetsche, B., Shalem, O., Wu, X., Makarova, K.S., et al. (2015). In vivo genome editing using Staphylococcus aureus Cas9. Nature 520, 186–191.

Sanjana, N.E., Shalem, O., and Zhang, F. (2014). Improved vectors and genome-wide libraries for CRISPR screening. Nature Methods 11, 783–784.

Sanson, K.R., Hanna, R.E., Hegde, M., Donovan, K.F., Strand, C., Sullender, M.E., Vaimberg, E.W., Goodale, A., Root, D.E., Piccioni, F., et al. (2018). Optimized libraries for CRISPR-Cas9 genetic screens with multiple modalities. Nature Communications 9.

Shalem, O., Sanjana, N.E., Hartenian, E., Shi, X., Scott, D.A., Mikkelsen, T.S., Heckl, D., Ebert, B.L., Root, D.E., Doench, J.G., et al. (2014). Genome-Scale CRISPR-Cas9 Knockout Screening in Human Cells. Science 343, 84–87.

Wang, J., Meng, X., Hu, X., Sun, T., Li, J., Wang, K., and Yu, H. (2019). XC as9 expands the scope of genome editing with reduced efficiency in rice. Plant Biotechnology Journal 17, 709–711.

Wu, X., Scott, D.A., Kriz, A.J., Chiu, A.C., Hsu, P.D., Dadon, D.B., Cheng, A.W., Trevino, A.E., Konermann, S., Chen, S., et al. (2014). Genome-wide binding of the CRISPR endonuclease Cas9 in mammalian cells. Nature Biotechnology 32, 670–676.

Xie, S., Cooley, A., Armendariz, D., Zhou, P., and Hon, G.C. (2018). Frequent sgRNA-barcode recombination in single-cell perturbation assays. PLOS ONE 13, e0198635.

Zhang, Y., Ge, X., Yang, F., Zhang, L., Zheng, J., Tan, X., Jin, Z.-B., Qu, J., and Gu, F. (2015). Comparison of non-canonical PAMs for CRISPR/Cas9-mediated DNA cleavage in human cells. Scientific Reports 4.

Zhong, Z., Sretenovic, S., Ren, Q., Yang, L., Bao, Y., Qi, C., Yuan, M., He, Y., Liu, S., Liu, X., et al. (2019). Improving Plant Genome Editing with High-Fidelity xCas9 and Non-canonical PAM-Targeting Cas9-NG. Molecular Plant 12, 1027–1036.

