## Supplementary Figures and Tables for "High-throughput screens of PAM-flexible Cas9 variants for gene knock-out and transcriptional modulation"

**Table S1. Composition of CRISPR libraries for each gene.**

| Gene | Region | PAM | No of sgRNAs |
| --- | --- | --- | --- |
| CD45 | CDS | NGG | 53 |
|  |  | NGA | 89 |
|  |  | NGC | 55 |
|  |  | NGT | 87 |
|  |  | NAN | 150 |
|  |  | NCN | 150 |
|  |  | NTN | 150 |
| | TSS ($\pm 1.5$ kb) | NGG | 123 |
|  |  | NGA | 211 |
|  |  | NGC | 135 |
|  |  | NGT | 206 |
|  |  | NAN | 150 |
|  |  | NCN | 150 |
|  |  | NTN | 150 |
|  | Non targeting |  | 250 |

| Gene | Region | PAM | No of sgRNAs |
| --- | --- | --- | --- |
| CD46 | CDS | NGG | 62 |
|  |  | NGA | 72 |
|  |  | NGC | 35 |
|  |  | NGT | 80 |
|  |  | NAN | 150 |
|  |  | NCN | 150 |
|  |  | NTN | 150 |
| | TSS ($\pm 1.5$ kb) | NGG | 393 |
|  |  | NGA | 274 |
|  |  | NGC | 366 |
|  |  | NGT | 227 |
|  |  | NAN | 150 |
|  |  | NCN | 150 |
|  |  | NTN | 150 |
|  | Non targeting |  | 250 |

| Gene | Region | PAM | No of sgRNAs |
| --- | --- | --- | --- |
| CD55 | CDS | NGG | 55 |
|  |  | NGA | 70 |
|  |  | NGC | 31 |
|  |  | NGT | 68 |
|  |  | NAN | 150 |
|  |  | NCN | 150 |
|  |  | NTN | 150 |
| | TSS ($\pm 1.5$ kb) | NGG | 400 |
|  |  | NGA | 280 |
|  |  | NGC | 341 |
|  |  | NGT | 300 |
|  |  | NAN | 150 |
|  |  | NCN | 150 |
|  |  | NTN | 150 |
|  | Non targeting |  | 250 |

**Table S2. sgRNA sequences for arrayed validation.**

| sgRNA ID | Target | PAM | Sequence |
| --- | --- | --- | --- |
| CD46_NGG_1 | CD46 exon 2 | TGG | CCGATCACAAATAGTATGGG |
| CD46_NGG_2 | CD46 exon 2 | AGG | CAAATAGTATGGGTGGCAAG |
| CD46_NGG_3 | CD46 exon 2 | AGG | ATAGTATGGGTGGCAAGAGG |
| CD46_NGG_4 | CD46 exon 2 | TGG | TTTGTGATCGGAATCATACA |
| CD46_NGG_5 | CD46 exon 2 | TGG | TCCATAGCTTCAAATGTTGG |
| CD46_NGG_6 | CD46 exon 3 | TGG | TCCCATTTGCAGGGACTGCT |
| CD46_NGG_7 | CD46 exon 3 | TGG | GCAAATGGGACTTACGAGTT |
| CD46_NGG_8 | CD46 exon 3 | AGG | AACTCGTAAGTCCCATTTGC |
| CD46_NGG_9 | CD46 exon 3 | TGG | GGCCAAGCAGTCCCTGCAAA |
| CD46_NGG_10 | CD46 exon 3 | GGG | ACTCGTAAGTCCCATTTGCA |
| CD46_NGG_11 | CD46 exon 3 | GGG | GCCAAGCAGTCCCTGCAAAT |
| CD46_GGN_1 | CD46 exon 2 | GGC | CGATCACAAATAGTATGGGT |
| CD46_GGN_2 | CD46 exon 2 | GGA | AAATAGTATGGGTGGCAAGA |
| CD46_GGN_3 | CD46 exon 2 | GGT | TAGTATGGGTGGCAAGAGGA |
| CD46_GGN_4 | CD46 exon 2 | GGC | TTGTGATCGGAATCATACAT |
| CD46_GGN_5 | CD46 exon 2 | GGC | CCATAGCTTCAAATGTTGGT |
| CD46_GGN_6 | CD46 exon 3 | GGC | CCCATTTGCAGGGACTGCTT |
| CD46_GGN_7 | CD46 exon 3 | GGT | CAAATGGGACTTACGAGTTT |
| CD45_NGG_1 | CD45 TSS | CGG | CTAGGTGATGATGTCAGATT |
| CD45_NGG_2 | CD45 TSS | TGG | CAGTTCATGCAGCTAGCAAG |
| CD45_NGA_1 | CD45 TSS | GGA | CTGTAAGGGTCCTCTTTGCA |
| CD45_NGA_2 | CD45 TSS | AGA | AACTGCTAGGTGATGATGTC |
| CD45_NGA_3 | CD45 TSS | TGA | TAGCTGCATGAACTGCTAGG |
| CD45_NGT_1 | CD45 TSS | AGT | CCTGCAAAGAGGACCCTTAC |
| CD45_NGT_2 | CD45 TSS | GGT | AGTTCATGCAGCTAGCAAGT |
| CD45_NGT_3 | CD45 TSS | TGT | CATGCAGCTAGCAAGTGGTT |
| CD45_NGC_1 | CD45 TSS | TGC | AATACTGTAAGGGTCCTCTT |
| CD45_NGC_2 | CD45 TSS | AGC | CGAATCTGACATCATCACCT |
| CD45_NGC_3 | CD45 TSS | AGC | CTAAGAACAAACCACTTGCT |
| CD45_NAG_1 | CD45 TSS | CAG | TCCTGCAAAGAGGACCCTTA |
| CD45_NAG_2 | CD45 TSS | AAG | GTAAGGGTCCTCTTTGCAGG |
| CD45_NAG_3 | CD45 TSS | CAG | AATCTGACATCATCACCTAG |
| CD45_NAA_1 | CD45 TSS | GAA | TGTAAGGGTCCTCTTTGCAG |
| CD45_NAA_2 | CD45 TSS | AAA | GGTTTTACTAACTTCTCCAA |
| CD45_NAA_3 | CD45 TSS | CAA | CTAGCAGTTCATGCAGCTAG |
| CD45_NTG_1 | CD45 TSS | TTG | AAATACTGTAAGGGTCCTCT |
| CD45_NTG_2 | CD45 TSS | ATG | GCTGCATGAACTGCTAGGTG |
| CD45_NTG_3 | CD45 TSS | TTG | TCATGCAGCTAGCAAGTGGT |
| CD45_Nontargeting_1 | Non-targeting | - | GTAGGCGCGCCGCTCTCTAC |
| CD45_Nontargeting_2 | Non-targeting | - | TACTAACGCCGCTCCTACAG |
| CD45_Nontargeting_3 | Non-targeting | - | ACGGAGGCTAAGCGTCGCAA |

**Table S3. PCR primer sequences.**

Stagger (between 1 and 9 nucleotides) is shown in brackets; examples of 8 nucleotide i7 and i5 barcodes for sample multiplexing are shown in **bold**. A full set of PCR2 primers (with multiple barcodes and stagger) is available for download here (readout primers v. July 2014): http://sanjanalab.org/lib.html

| Function | Sequence |
| --- | --- |
| PCR1, amplify sgRNA cassette for CRISPR library sequencing (F) | TCTTGTGGAAAGGACGAAACACCG |
| PCR1, amplify sgRNA cassette for CRISPR library sequencing (R) | CCGACTCGGTGCCACTTTTTCAA |
| PCR2, add adaptors for CRISPR/indel library sequencing (F) | AATGATACGGCGACCACCGAGATCTACACTCTTTCCCTACACGACGCTCTTCCGATCT (ACGATCGAT) **AGGTAAGG**TCTTGTGGAAAGGACGAAACACCG |
| PCR2, add adaptors for CRISPR/indel library sequencing (R) | CAAGCAGAAGACGGCATACGAGAT**AGGTAAGG**GTGACTGGAGTTCAGACGTGTGCTCTTCCGATCT (ACGATCGAT) CCGACTCGGTGCCACTTTTTCAA |
| PCR1, amplify CD46 exon 2 indel (F) | TCTTGTGGAAAGGACGAAACACCGTACCTGCTGCCAGACCACAG |
| PCR1, amplify CD46 exon 2 indel (R) | CCGACTCGGTGCCACTTTTTCAAGTATGGGTGGCAAGAGGAGG |
| PCR1, amplify CD46 exon 2 indel (F) | TCTTGTGGAAAGGACGAAACACCGGAAGCTATGGAGCTCATTGG |
| PCR1, amplify CD46 exon 2 indel (R) | CCGACTCGGTGCCACTTTTTCAAAAGAGGTTTGTCTTACTTACTATAACAG |
| PCR1, amplify CD46 exon 3 indel (F) | TCTTGTGGAAAGGACGAAACACCGCTTTCAGGAGAAACATGTCC |
| PCR1, amplify CD46 exon 3 indel (R) | CCGACTCGGTGCCACTTTTTCAATTACTAACACTTCCCTTATTTCC |

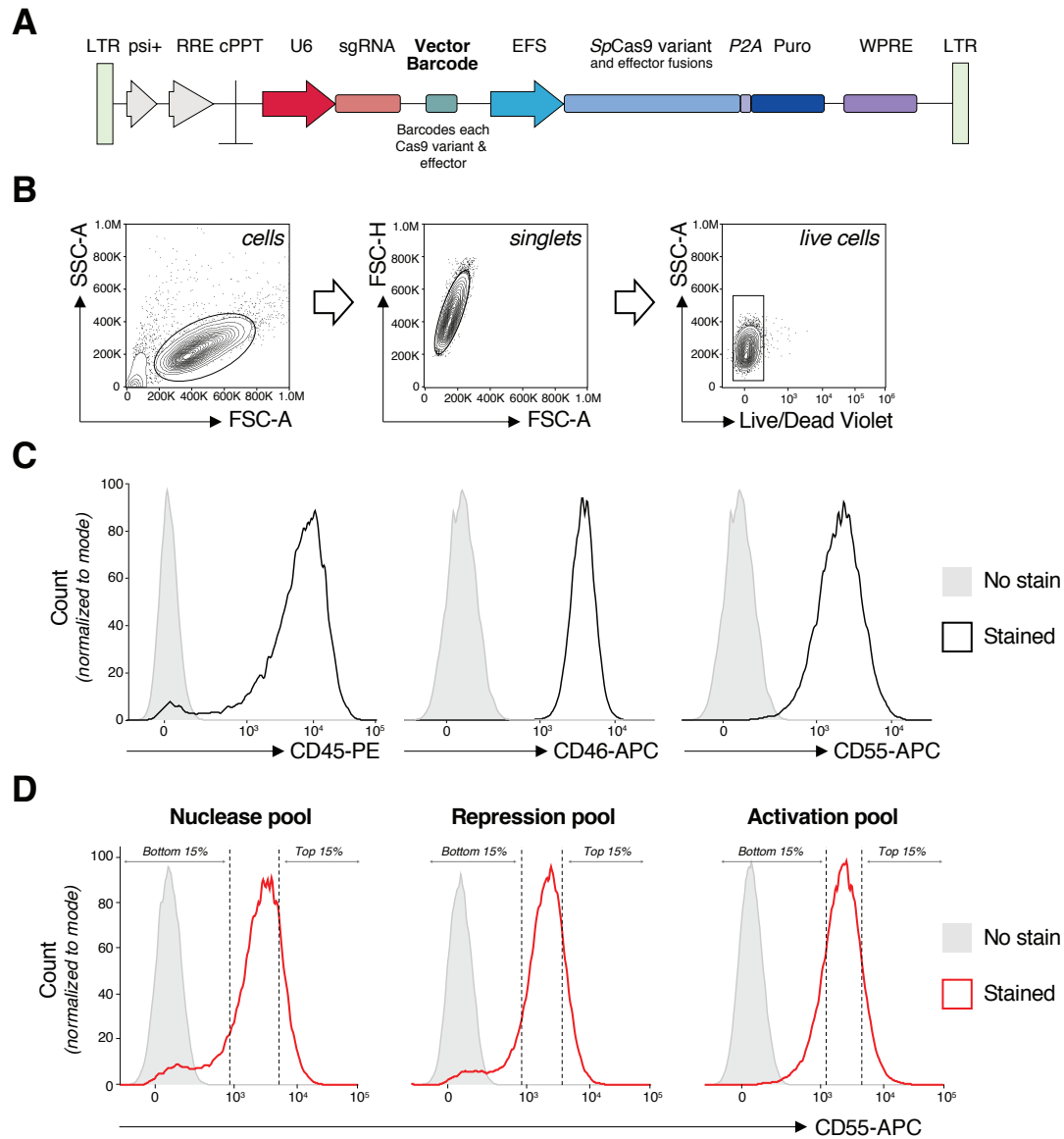

**Figure S1. Lentiviral vector design and screen gating strategy.** **A)** Lentiviral expression vector for Cas9 and sgRNA expression, based on the lentiCRISPRv2 system (Sanjana et al., 2014). sgRNA scaffold was modified for improved activity (Chen et al, 2013)<sup>29</sup>. A six nucleotide barcode was inserted between the sgRNA scaffold and the EFS promoter to enable identification of Cas9 variant and effector. For transcriptional repression, KRAB domain was fused to catalytically inactive Cas9 (dCas9, D10A, H840A) on N-terminus; for transcriptional activation, VP64-p65-Rta (VPR) complex was fused to catalytically inactive Cas9 (dCas9, D10A, H840A) on C-terminus. In all cases Cas9 sequences were codon optimized for human expression and contained a C-terminal nuclear localization signal (NLS). Long terminal repeat (LTR), psi packaging signal (psi+), rev response element (RRE), central polypurine tract (cPPT), elongation factor-1α short promoter (EFS), 2A self-cleaving peptide (P2A), puromycin selection marker (puro), posttranscriptional regulatory element (WPRE). **B)** Gating strategy for flow cytometry analyses and sorting. The cells were separated from debris based on forward (FSC) and side scatter (SSC) properties, followed by doublet and dead cell exclusion. **C)** Surface expression of CD45, CD46 and CD55 in K562 cell line. **D)** Sorting gates for CRISPR nuclease, repression and activation pools in K562 cells. Representative histograms for one of the genes (CD55) are shown.

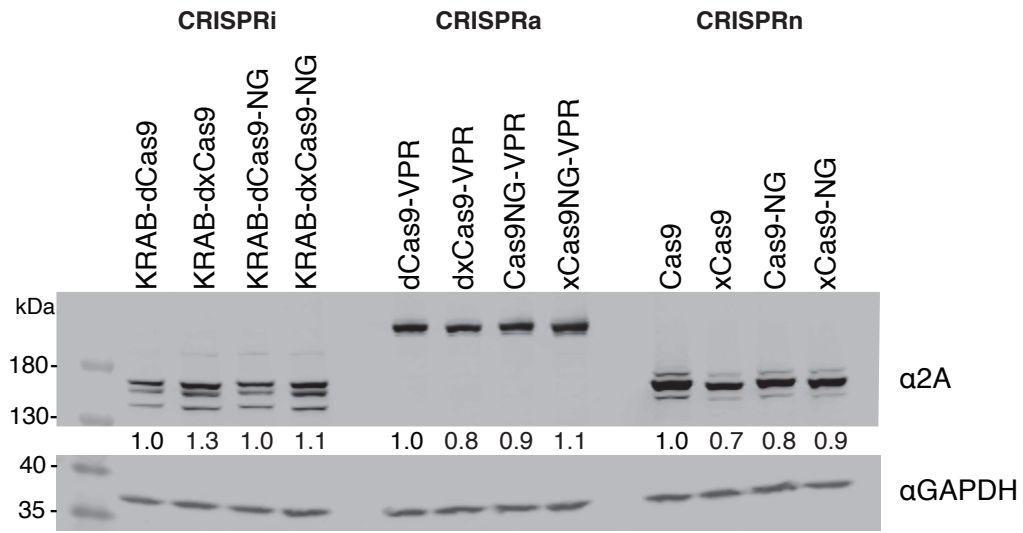

**Figure S2. Quantification of the expression levels of Cas9 variants.** Cas9 expression levels were quantified by western blot using an antibody against the 2A peptide found at the C-terminus of all variants. The expression of Cas9 variants was normalized by the expression level of GAPDH in the same sample. For all three types of Cas9 effectors (CRISPRi, CRISPRa and CRISPRn) the expression levels were normalized to the relative expression level of wild-type Cas9, which was set to 1.

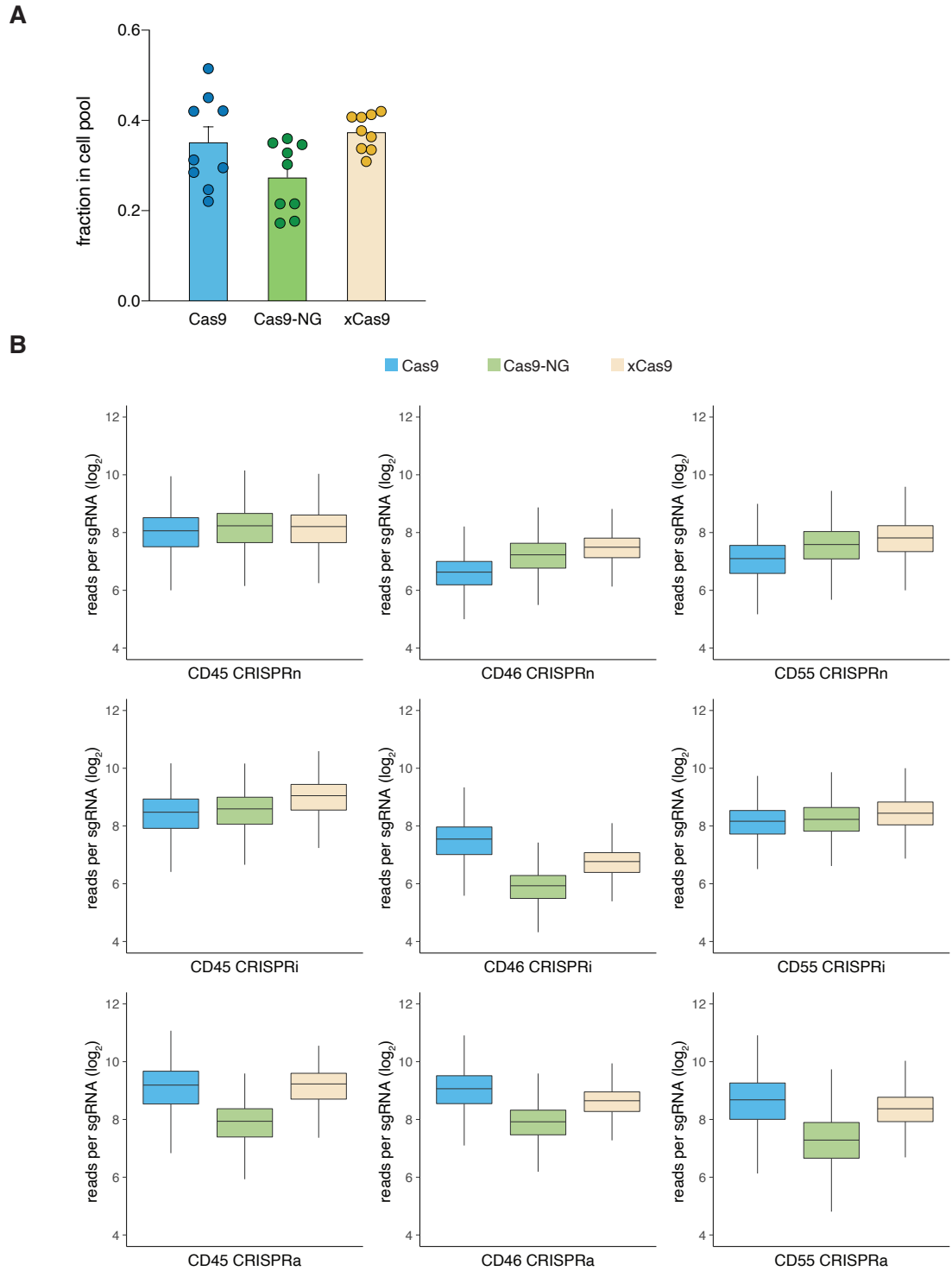

**Figure S3. Relative abundance of barcodes corresponding to each Cas9 effector in mixed population pre-sort.** A) Relative frequency of each Cas9 variant in each of the nine cell pools used for screening. Individual values and standard error of the mean are shown. B) Read count distributions for every sgRNA with each Cas9 effector shown separately for each screen pool.

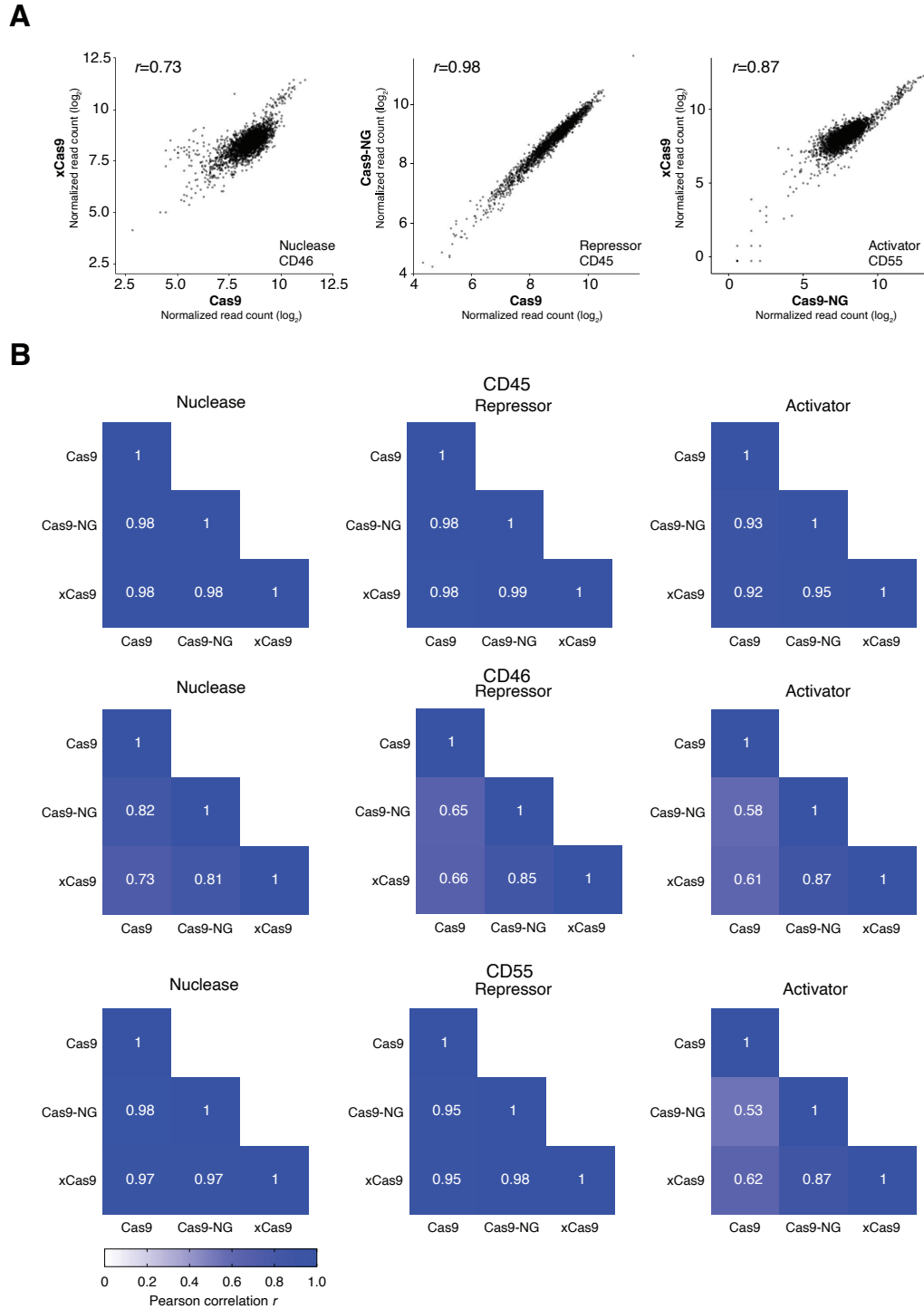

**Figure S4. sgRNA count correlations between Cas9 enzymes within the same pool before sorting. A)** Representative scatterplots for nuclease, repressor and activator pools. $r$ indicates Pearson correlation coefficient. **B)** Pearson correlation coefficient of normalized read counts for each Cas9 combination within the same pool (modality + gene).

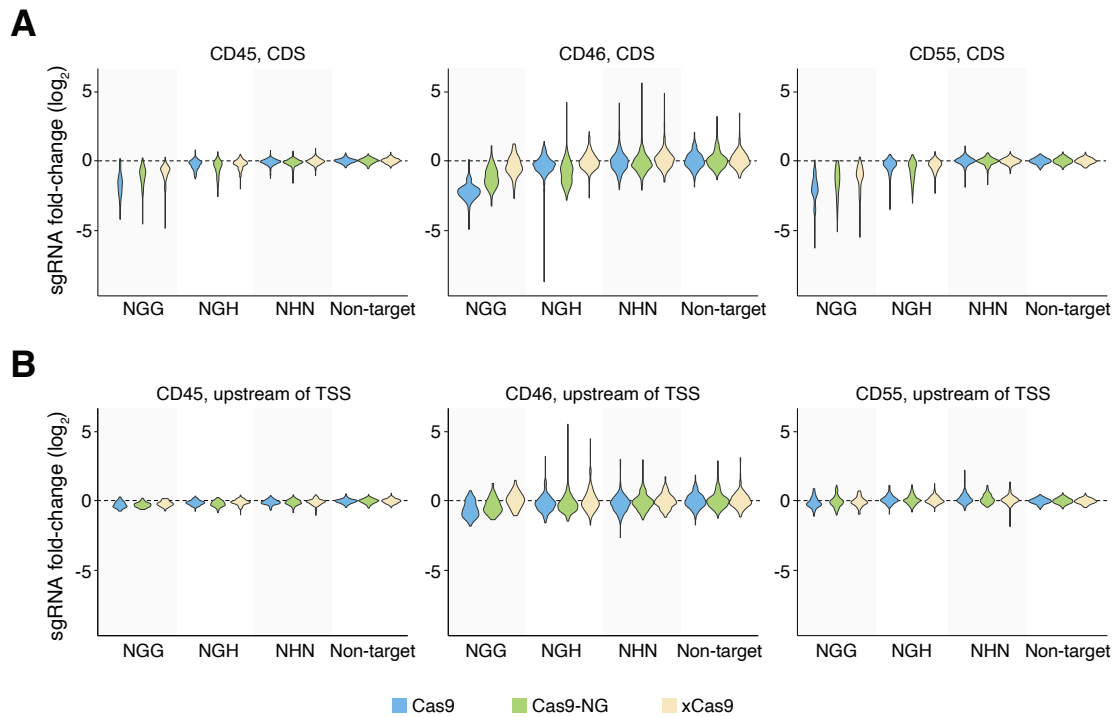

**Figure S5. Nuclease CRISPR competition screen for CD45, CD46, and CD55.** **A)** Fold-change of CDS-targeting sgRNAs for each Cas9 variant in cell populations expressing high levels of target gene (CD45: $n = 1,027$ sgRNAs; CD46: $n = 1,025$ sgRNAs; CD55: $n = 1,122$ sgRNAs) over low-expressing cells is shown. **B)** Fold-change of sgRNAs targeting between 1.5 kb and 0.5 kb upstream of the TSS in cell populations expressing high level of target gene (CD45: $n = 655$ sgRNAs; CD46: $n = 725$ sgRNAs; CD55: $n = 781$ sgRNAs) over low-expressing cells is shown. Median of non-targeting (NT) sgRNAs was used to normalize fold-change for each enzyme.

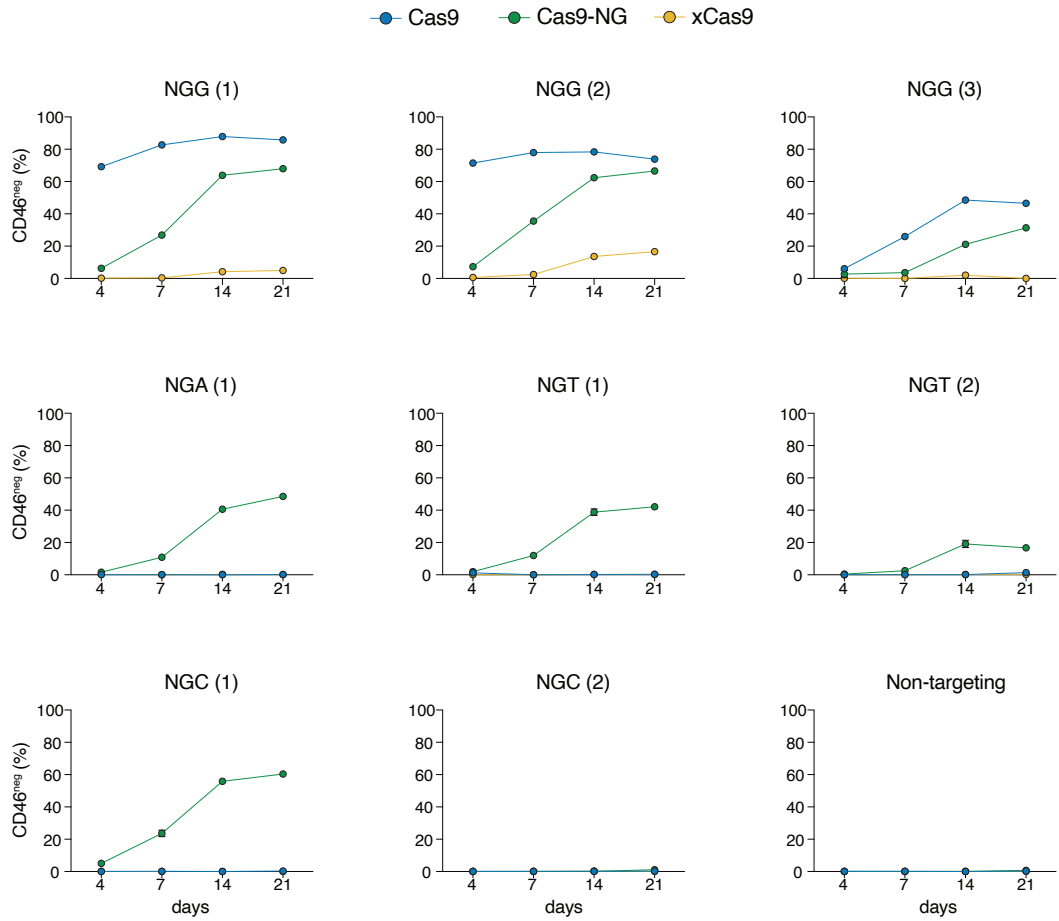

**Figure S6. Time course of CD46 knock-out by Cas9 variant.** CD46<sup>+</sup> A375 cell line was transduced with lentivirus encoding each of three Cas9 variants and sgRNAs targeting CD46 coding sequences with the indicated PAMs. Following selection, CD46 negative cells were quantified based on the gate set on the unstained population. Standard error of the mean ($n = 3$ replicate transductions) is shown.

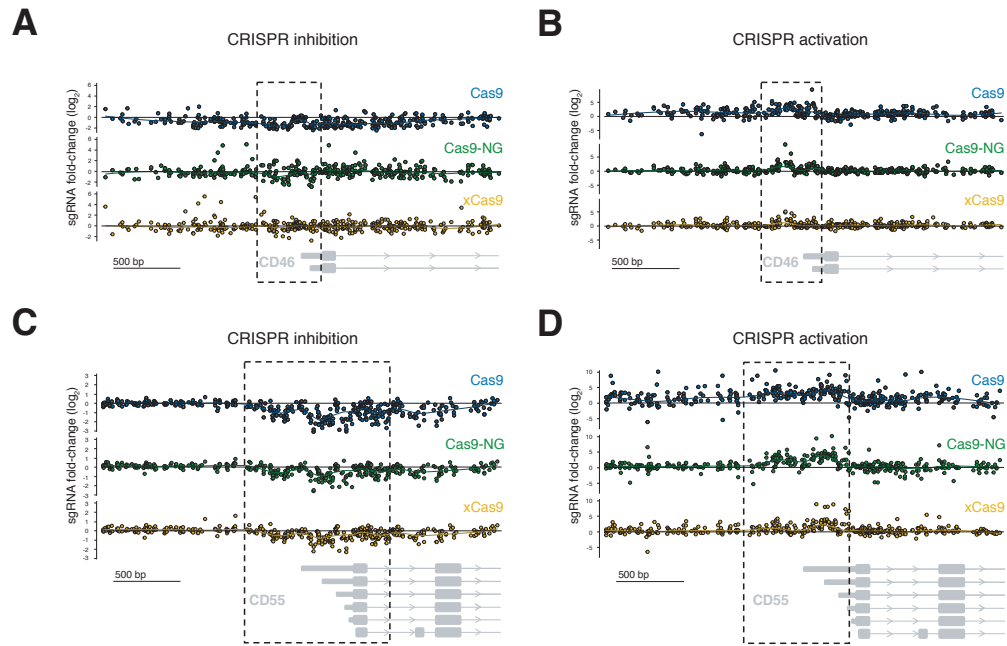

**Figure S7. High-throughput CRISPR transcriptional modulation screen at CD46 and CD55.** Fold-change of sgRNAs targeting the 3 kb region surrounding the primary TSS of CD46 (A and B) and CD55 (C and D) genes. Only sgRNAs associated with the NGG PAM are displayed (CD46: $n = 393$ sgRNAs; CD55: $n = 400$ sgRNAs). The regions with strongest NGG sgRNA activity (indicated with dashed lines) were used to select sgRNAs for subsequent analyses. Collapsed gene models of CD46 and CD55 isoforms (Ensembl CD46-201, CD46-206; CD55-204, CD55-203, CD55-206, CD55-201, CD55-213, CD55-202) are shown in grey.

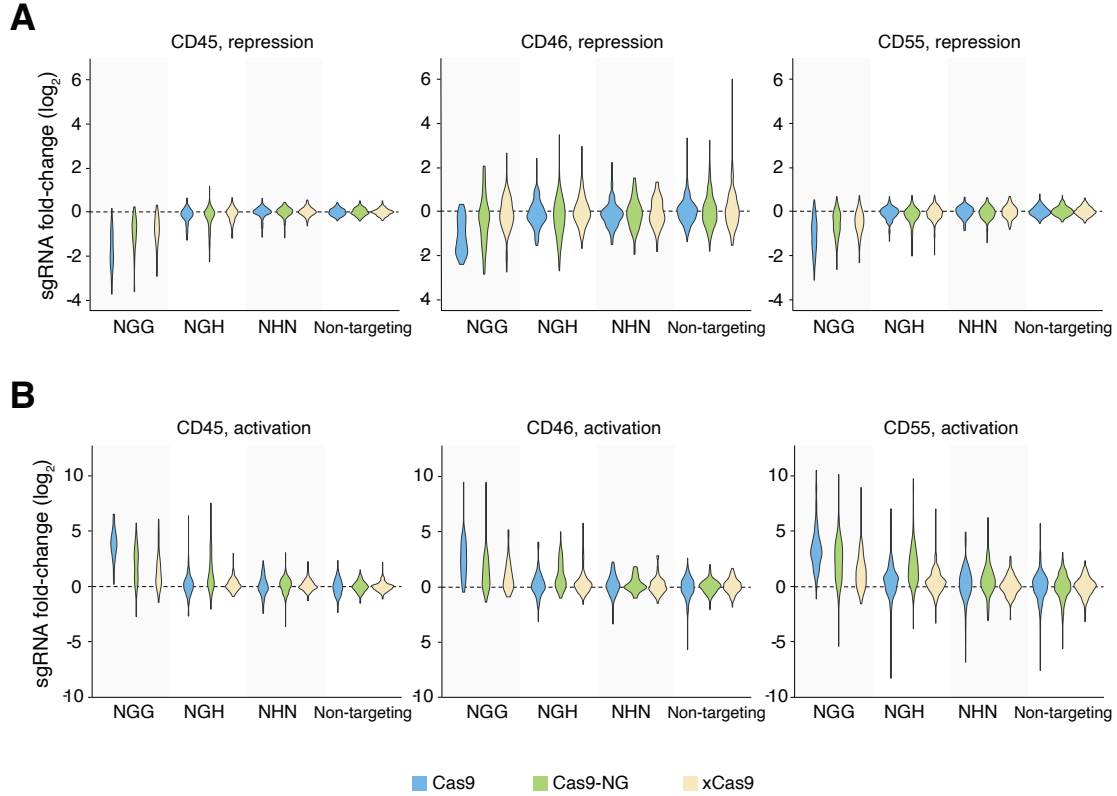

**Figure S8. Transcriptional modulation CRISPR competition screen results for sgRNAs targeting the optimal region surrounding the TSS.** Fold-change of sgRNAs in sorted cell populations expressing high level of target gene over low-expressing cells is shown (**A** - CRISPR inhibition, **B** - CRISPR activation). The median of the non-targeting sgRNAs was used to normalize fold-change for each Cas9 variant. Only sgRNAs targeting the optimal region surrounding the gene TSS are included in the analysis (CRISPR inhibition, CD45: $n = 624$ sgRNAs; CD46: $n = 577$ sgRNAs; CD55: $n = 946$ sgRNAs; CRISPR activation, CD45: $n = 670$ sgRNAs; CD46: $n = 506$ sgRNAs; CD55: $n = 804$ sgRNAs).

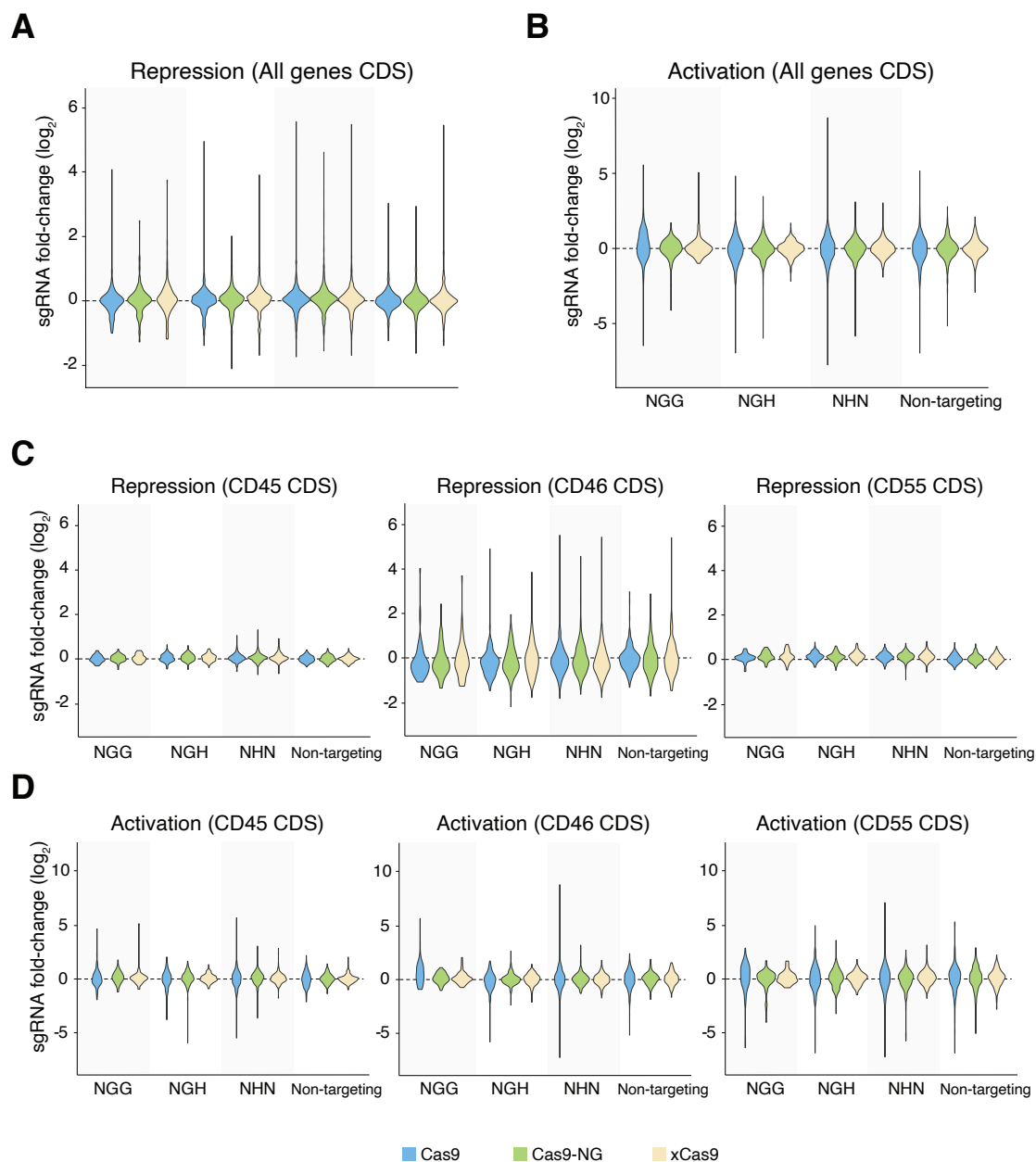

**Figure S9. Transcriptional modulation CRISPR competition screen results for CDS targeting sgRNAs.** Fold-change of sgRNAs in cell populations expressing high level of target gene over low-expressing cells is shown. Median of non-targeting (NT) sgRNAs was used to normalize fold-change for each enzyme. Only sgRNAs targeting CDS exons are included in the analysis. **A)** CRISPR inhibition screen and **B)** CRISPR activation screen – all three genes together. **C)** CRISPR inhibition screen and **D)** CRISPR activation screen split by gene: CD45: $n = 984$ sgRNAs; CD46: $n = 949$ sgRNAs; CD55: $n = 924$ sgRNAs

**A**

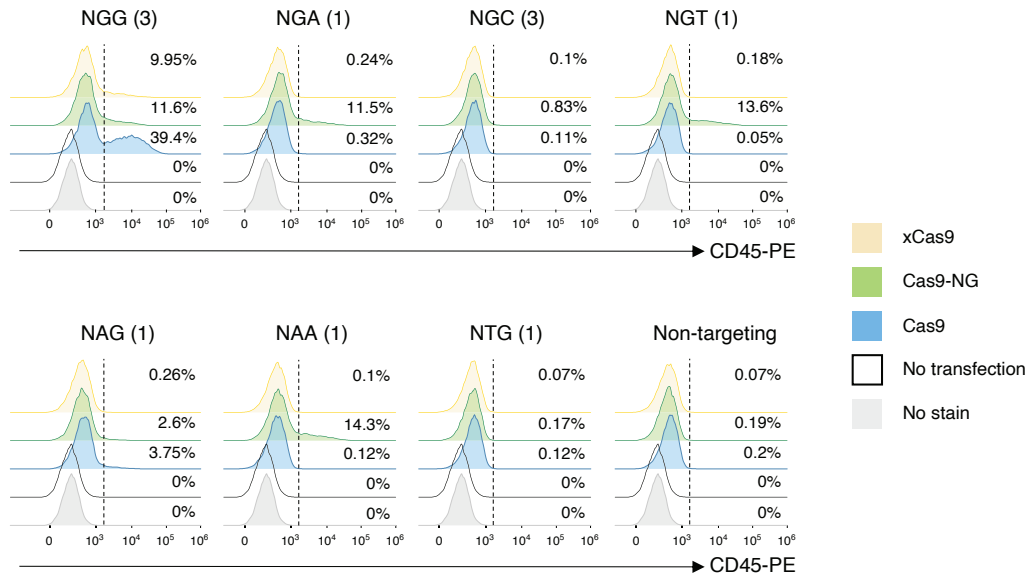

**B**

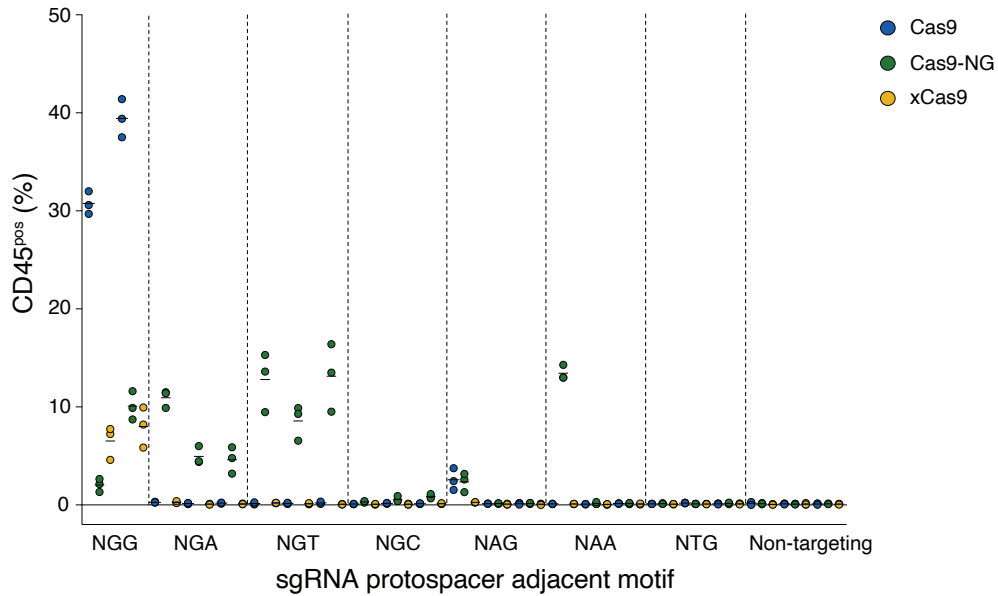

**Figure S10. CD45 expression following CRISPR activation in CD45<sup>neg</sup> A375 cell line. A)** Representative histograms of CD45 staining for each PAM category. Dashed line indicates CD45<sup>pos</sup> gate; numbers on histograms correspond to the percentage of cells in CD45<sup>pos</sup> gate. **B)** CD45 positive cells after CRISPRa transduction. Mean and individual values from three independent experiments are shown ($n = 23$ sgRNAs, each tested with all three Cas9 variants).

**A**

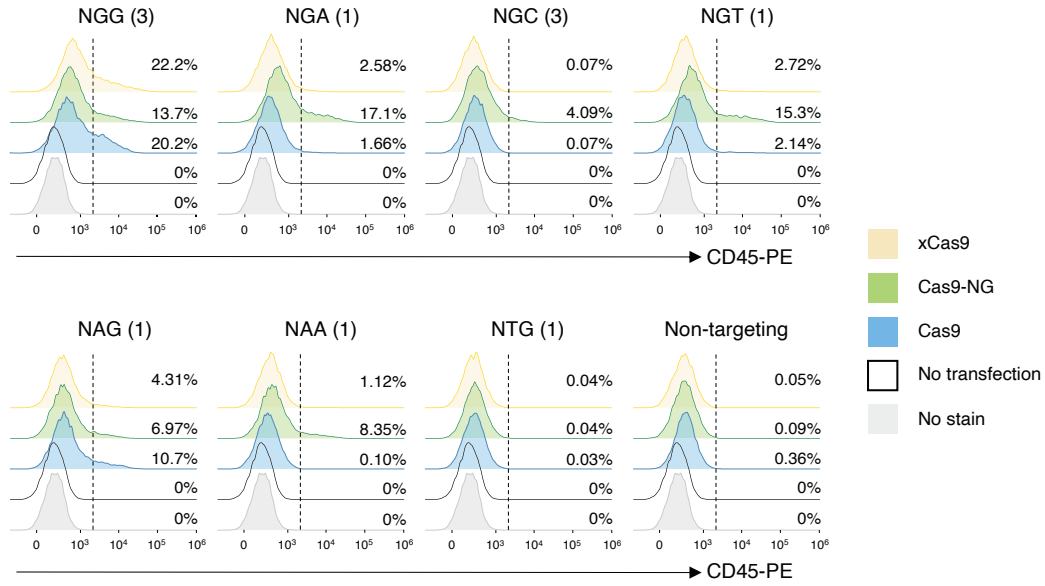

**B**

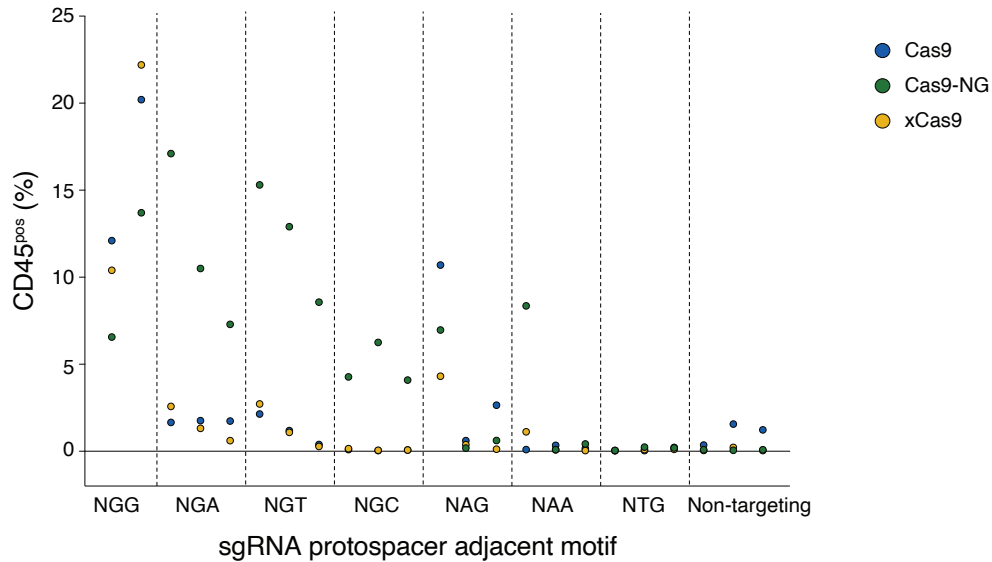

**Figure S11. CD45 expression following CRISPR activation in CD45<sup>neg</sup> HEK-293FT cell line. A)** Representative histograms of CD45 staining for each PAM category. Dashed line indicates CD45<sup>pos</sup> gate; numbers on histograms correspond to the percentage of cells in CD45<sup>pos</sup> gate. **B)** CD45 positive cells after CRISPRa transduction. Two (NGG) or three (all other PAM categories) sgRNAs per PAM were tested in a single experiment. $n = 23$ sgRNAs, each tested with all three Cas9 variants.

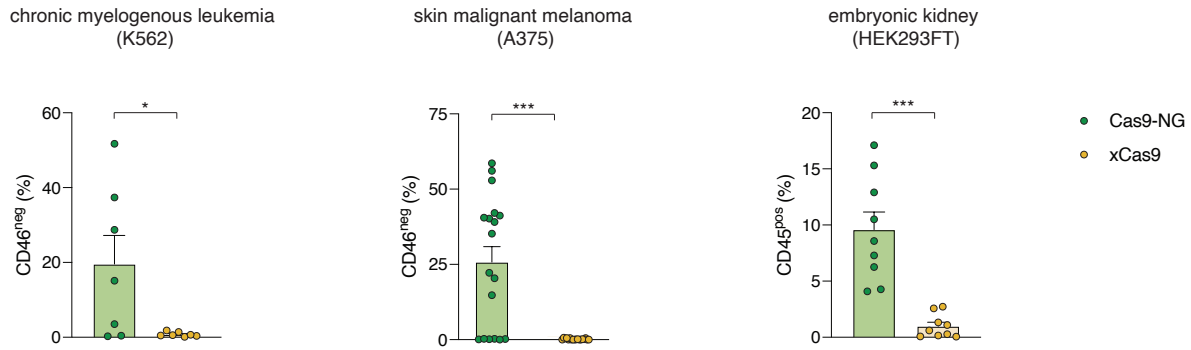

**Figure S12. Cas9-NG outperforms xCas9 at NGH PAM sites across different cell lines.** Frequency of cells with modulated expression of target proteins was calculated based on gates set on the negative or positive populations, respectively. For K562 and A375, CRISPR nuclease activity on CD46 gene is shown. For HEK293FT, CRISPR activation of CD45/PTPRC gene is shown. Individual values and standard error of the mean are displayed. sgRNA sequences are listed in Table S2. Student's t-test, ns $p > 0.05$, \* $p < 0.05$, \*\*\* $p < 0.001$. The data (combined here for comparison purposes) are originally from Figures 1E, S6 and S14.

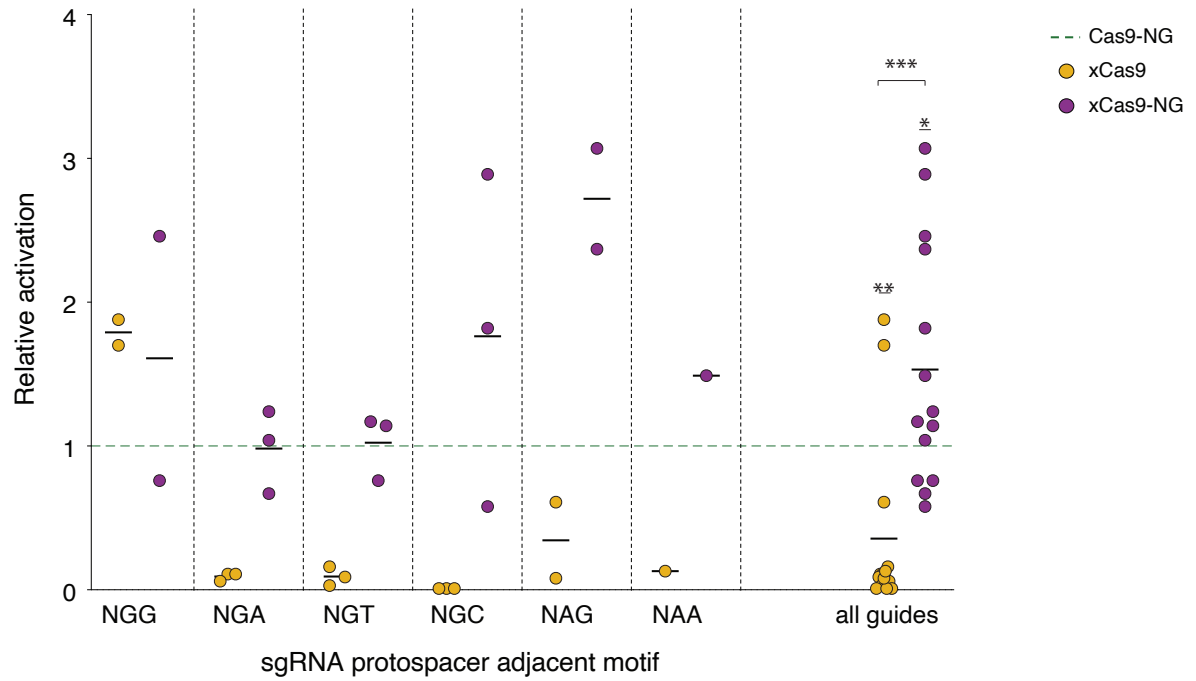

**Figure S13. Comparison of CD45 transcriptional activation efficiency between existing PAM-flexible Cas9 variants and xCas9-NG in HEK-293FT cell line.** Activation efficiency for xCas9 and xCas9-NG was normalized to Cas9-NG on a per-sgRNA basis. Individual values for each sgRNA are shown with a solid line indicated the mean over sgRNAs with the same PAM. Student's t-test results are shown between xCas9 and Cas9-NG (\*\* $p < 0.01$), xCas9-NG and Cas9-NG (\* $p < 0.05$) and xCas9-NG and xCas9 (\*\*\*) $p < 0.001$).

**A**

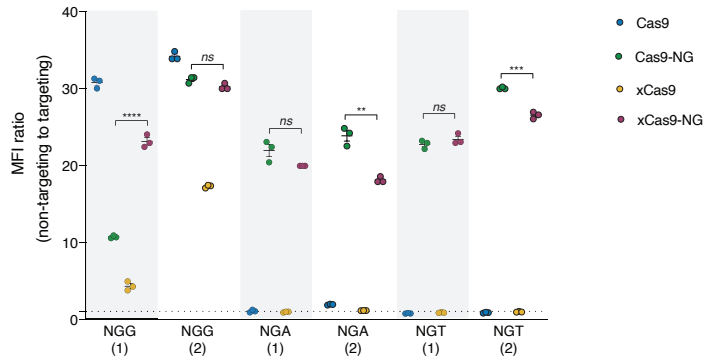

**B**

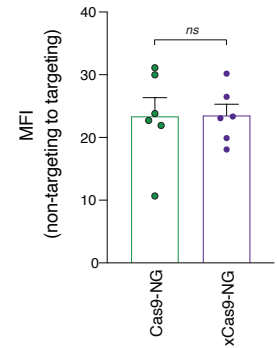

**Figure S14. CD45 expression following CRISPR inhibition in CD45<sup>+</sup> cell line K562.** **A)** Reduction of CD45 expression following transduction with Cas9 effectors and sgRNAs targeting the transcription start site of CD45/PTPRC gene (sgRNA sequences are shown in Table S2). For each Cas9 variant and sgRNA, the median fluorescence intensity (MFI) of CD45 staining was normalized by dividing the MFI of corresponding non-targeting sgRNAs by targeting sgRNAs. Individual values and standard error of the mean (3 replicates per sgRNA) are shown. **B)** Knock-down activity for individual sgRNAs with target sites with the indicated PAMs. For each sgRNA, mean of three replicates is shown. Error bars indicate standard error of the mean. Two-sided t-test: ns, $p > 0.05$, \*\* $p < 0.01$, \*\*\* $p < 0.001$, \*\*\*\* $p < 0.0001$.

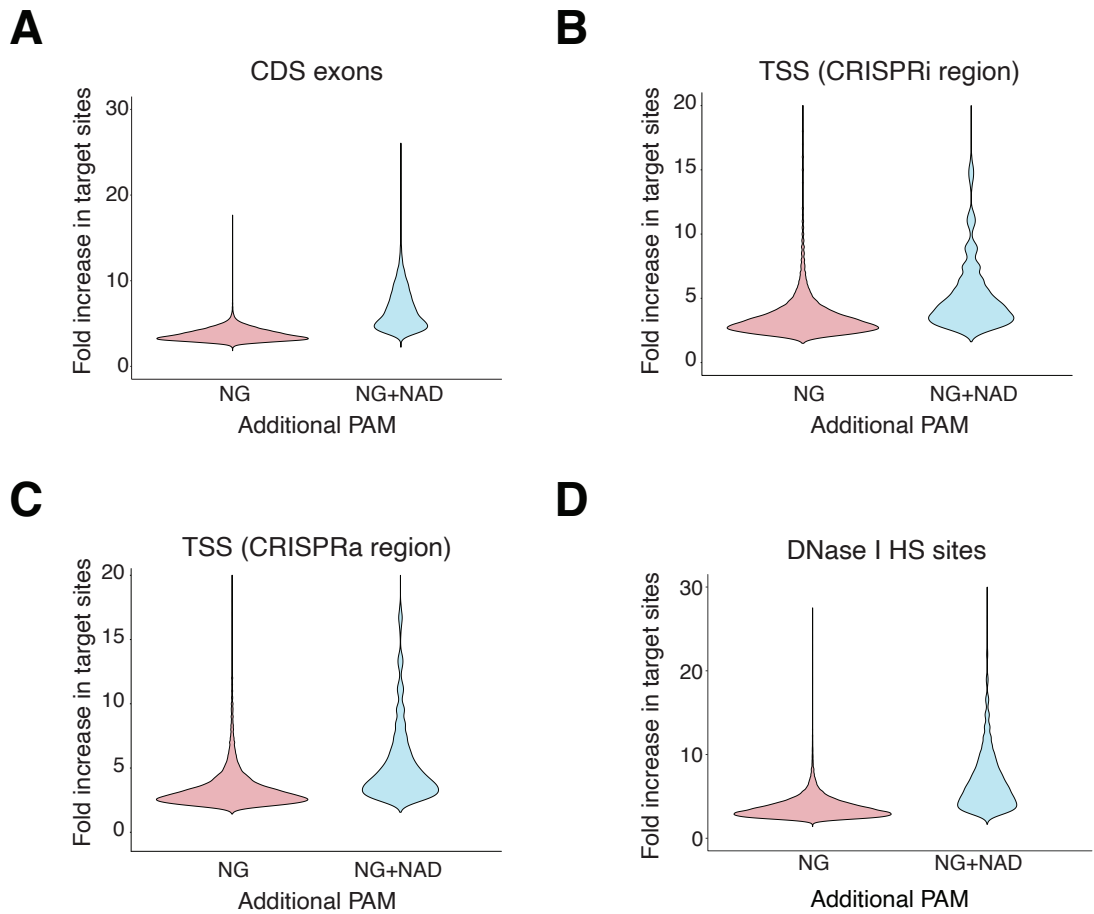

**Figure S15. Targetable sites with NG and NG+NAD PAMs in all protein-coding genes in the human genome.** Targetable sites when using additional PAMs (beyond NGG) in **A)** coding exons of 18,544 human protein-coding genes, **B)** near transcription start sites in optimal CRISPRi regions, **C)** near transcription start sites in optimal CRISPRa regions, and **D)** 202,000 DNase I hypersensitivity (HS) sites in K562 cell line. For example, in CDS exons, there is a 3.7x mean increase in target sites with NG PAMs and a 6.6x mean increase with NAD and NG PAMs. All increases are given as fold-change over targeting NGG PAM sites alone.
